# A brainstem integrator for self-localization and positional homeostasis

**DOI:** 10.1101/2021.11.26.468907

**Authors:** En Yang, Maarten F. Zwart, Mikail Rubinov, Benjamin James, Ziqiang Wei, Sujatha Narayan, Nikita Vladimirov, Brett D. Mensh, James E. Fitzgerald, Misha B. Ahrens

## Abstract

To accurately track self-location, animals need to integrate their movements through space. In amniotes, representations of self-location have been found in regions such as the hippocampus. It is unknown whether more ancient brain regions contain such representations and by which pathways they may drive locomotion. Fish displaced by water currents must prevent uncontrolled drift to potentially dangerous areas. We found that larval zebrafish track such movements and can later swim back to their earlier location. Whole-brain functional imaging revealed the circuit enabling this process of positional homeostasis. Position-encoding brainstem neurons integrate optic flow, then bias future swimming to correct for past displacements by modulating inferior olive and cerebellar activity. Manipulation of position-encoding or olivary neurons abolished positional homeostasis or evoked behavior as if animals had experienced positional shifts. These results reveal a multiregional hindbrain circuit in vertebrates for optic flow integration, memory of self-location, and its neural pathway to behavior.

## Introduction

Many animals keep track of where they are in their environment^1^. For example, when animals visit unknown, potentially dangerous areas, they need to efficiently return to a safe location. They also must be able to revisit food-rich areas and avoid foraging in food-poor areas^2^. These navigational abilities are thought to be supported by neural representations of space in what is called a ‘cognitive map’^3–6^, arising from diverse brain regions including mammalian hippocampus^6–9^, entorhinal cortex^10,11^, hippocampus’ homologue in fish^12–14^, central complex of *Drosophila melanogaster*^15,16^, and retrosplenial cortex^17^. Navigational systems in mammals include subcortical angular neural integrators containing head-direction cells^18,19^. However, positional representations of self-location have not been observed in vertebrate hindbrain.

Navigation relies upon multiple sensory cues, including sensations generated by self-movement^20^, visual landmarks^21^, magnetic poles^22^, hydrostatic pressure^23^, electric discharge^24^, or a combination^25,26^. An important cue is optic flow, because the speeds and directions of motion signals across the retina provide information about an animal’s motion through its environment^27–33^. Extracting self-velocity from optic flow and integrating it over time^34^ produces a representation of how much an animal has moved, which works for self-generated movements (e.g., swimming, walking, flying) and movements imposed by the environment (e.g., fish moved by water flow, or birds by wind). Ants^35^, mammals^36,37^, and fruit flies^38^ show behaviors based on path integration. In mammals, networks including grid cells are thought to mediate path integration^5,11,39,40^. However, no complete neural transformation from an animal’s translational velocity, to a representation of self-location, to behavioral change, has been identified.

Larval zebrafish must control their location in space because, for instance, water currents could sweep them into dangerous areas. Animals can turn and swim toward water flow via rheotaxis^41,42^ and the optomotor response^29–31,43,44^. Yet it is unknown whether they explicitly track their location over long timescales and use memorized positional information to return to their earlier location — a behavior we term ‘positional homeostasis’. Such capabilities could be ethologically critical because larval zebrafish do not swim continuously^45,46^ and can be moved by currents during rest. Here, we discovered a multiregional circuit in the vertebrate hindbrain, with potential homologues in mammals, that computes self-location and communicates it to olivocerebellar circuits to guide locomotion.

## Zebrafish perform positional homeostasis

When landmarks are not available, efficient positional homeostasis requires path integration: summing the direction and magnitude of instantaneous velocity to compute overall displacement over prolonged time periods and thus enable corrective swim maneuvers. To investigate path integration, we exposed fish to stochastic virtual currents (visual flow in a closed-loop, one-dimensional, virtual-reality [VR] environment; Fig. 1A). Fish can approximately steady their location in space while swimming only a fraction of the time (Fig. 1B; a swim bout lasts ~100-400 ms; gaps of multiple seconds occur; at most, fish swim about once per second). To test if fish use an internal representation of self-location, we simulated positional displacements by varying sequences of virtual water currents in VR^47^ environments with fine stripes or random visual patterns. Fish swam in closed-loop for 10 seconds to determine their own starting location (Fig. 1C). Next, their position was changed in a *displace* period to a location determined by one of several pre-programmed trajectories that the fish could not control (open-loop^48^; displaced by multiple times their body length [~4 mm]). The trajectories were chosen so that fish would swim little or not at all, by using only short periods of translation, or limiting how much the fish was pulled backward (which triggers earlier responses than forward push). A *zero-displacement* trajectory was the baseline comparison. Fish were then held at the same location for several seconds in a *hold* period (open loop). In the subsequent *swim* period, fish were exposed to a virtual backward current and free to swim forward (closed-loop). The balance between the backward current pull and the amount of forward swimming during this period determined the fish’s final position, which is the starting position of the next trial. The safety of being in control may give that location positive ‘valence’^49^, motivating the fish to return there after the next trial’s forced displacement, which may have negative valence^48^.

**Figure 1:**
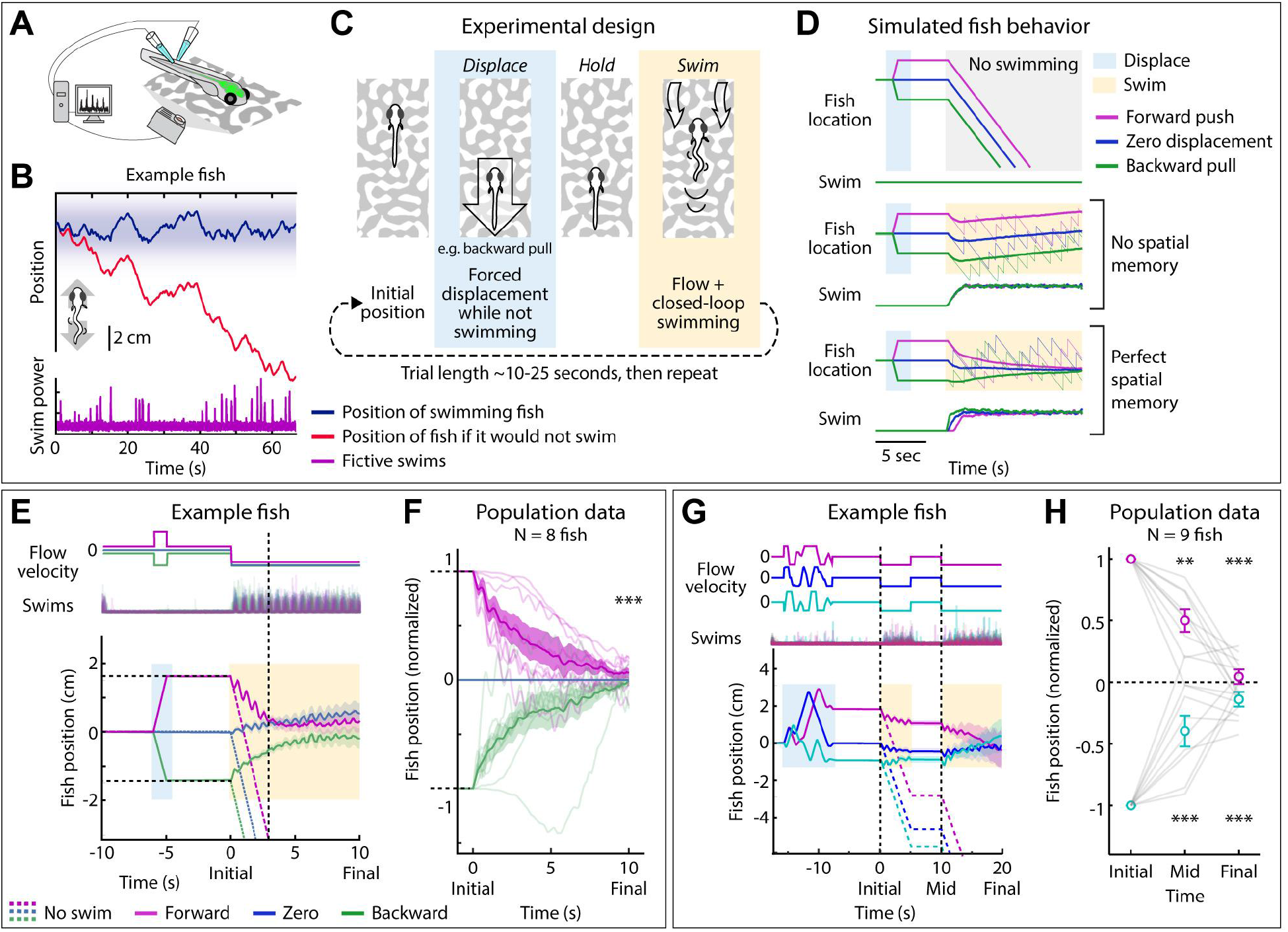
Larval zebrafish track their spatial location and correct for unintended displacements. (A) Experimental setup: larval zebrafish fictively swimming in a virtual reality environment. (B) Positional homeostasis: example fish stabilizes its location in stochastic virtual water current. (C) Experimental design: fish undergo one of various forced displacements (*displace*). After a period of no motion (*hold*), they correct for this displacement by swimming the correct amount against a virtual flow (*swim*). (Fish icons are enlarged and not to scale.) (D) Simulated fish behavior showing two classes of outcomes. If average trajectories during forward-push vs. backward-pull trial types converge during *swim*, it implies fish have memory of self-location. (E) Example fish in forward-push vs. backward-pull experiment. *Dashed lines*: trajectory that fish would take without swimming, to show speed of virtual current. *Solid lines*: actual trajectories for three trial types, which approximately converge at ~5s, showing this fish has memory of earlier displacement. (F) Trajectories (8 fish) normalized to zero-displacement trajectory converge at ~10s, indicating accurate memory of previous location shift. (One sample t-test for final positions, *** p<0.001, p=7.3e-8 for backward pull, specified mean=-1, p=6.3e-7 for forward push, specific mean=1.) (G) Assay to test integration during stochastic backward and forward motion, and to examine corrections made over two *swim* periods separated by pause. Example fish successfully corrects for stochastic displacements; correction continues across two *swim* periods. (H) Population data showing accurate correction distributed over both *swim* periods, i.e. path integration of complex trajectories. (One sample t-test, ** p<0.01, p=0.0023 for forward push at mid time point, *** p<0.001, p=2.5e-5 for backward pull at mid time point. p=2.1e-7 for forward push at final time point, p=5.8e-8 for backward pull at final time point.)

We asked whether fish compensated for displacement by analyzing if the swim-period trajectory converged with the *zero-displacement* trajectory or whether it remained separated by the amount of earlier passive displacement. A simulated fish that does not swim would be pulled backward by the current (Fig. 1D, *top*); a simulated memoryless fish that only responds to visual flow would swim but average trajectories for *forward-push, zero-displacement*, and *backward-pull* trials would remain separated (Fig. 1D, *middle*); while a fish with perfect spatial memory (simulated with a type of PID controller^50^; Methods) would respond to flow but also swim to cancel earlier displacements, so average trajectories for *forward, zero*, and *backward* trials would converge (Fig. 1D, *bottom*;).

In real fish, swim trajectories approximately converged: following net forward or net backward forced displacement, on average fish reached locations consistent with *zero-displacement* trajectories after about 10 seconds of swimming (Fig. 1E,F). This implies that fish retain an accurate memory of the forced displacement for up to 15 seconds.

Fish also corrected for more complex forced movements, such as multiple shifts or shifts of different durations (Suppl. Fig. 1A-D), whether or not they swam during the *displace* and *hold* periods. They also corrected for semi-random forward and backward trajectories followed by a *swim* period that was broken in two (Fig. 1G,H), so that the convergence continued over 20 seconds. Behavioral effects induced by displacement history included modulation of response time and swim vigor in the *swim* period (Suppl. Fig. 1E-G). Trajectories also converged following a change in motosensory gain, showing that fish use visual flow for positional homeostasis even when swim outcomes (amount of translation per unit of swimming) are altered, consistent with visually driven path integration (Suppl. Fig. 1H-J). Although slow forward or backward positional drift occurred during the closed-loop period in all trial types (Fig. 1E, Suppl. Fig. 2), likely reflecting error accumulation common to integrator circuits seen in other species^51^, the convergence of the three average trajectories implies that fish possess a persistent memory of location changes. Thus, fish integrate visual flow into a representation of location change and correct for unintended location changes by altered swimming. Location memory can persist for over 20 seconds.

## Zebrafish brainstem codes for self-location

To identify brain regions encoding self-location, we used whole-brain, light-sheet microscopy^52–54^ to record single-neuron activity using genetically encoded calcium indicators (Fig. 2B; Suppl. Fig. 3). To search for activity encoding displacements that occurred many seconds ago, we exposed fish to a forward push or backward pull followed by a 17-second pause. We found neurons that encoded the positional shift for the entire pause (Fig. 2C; Methods), showing that the brain persistently encodes past displacements.

**Figure 2:**
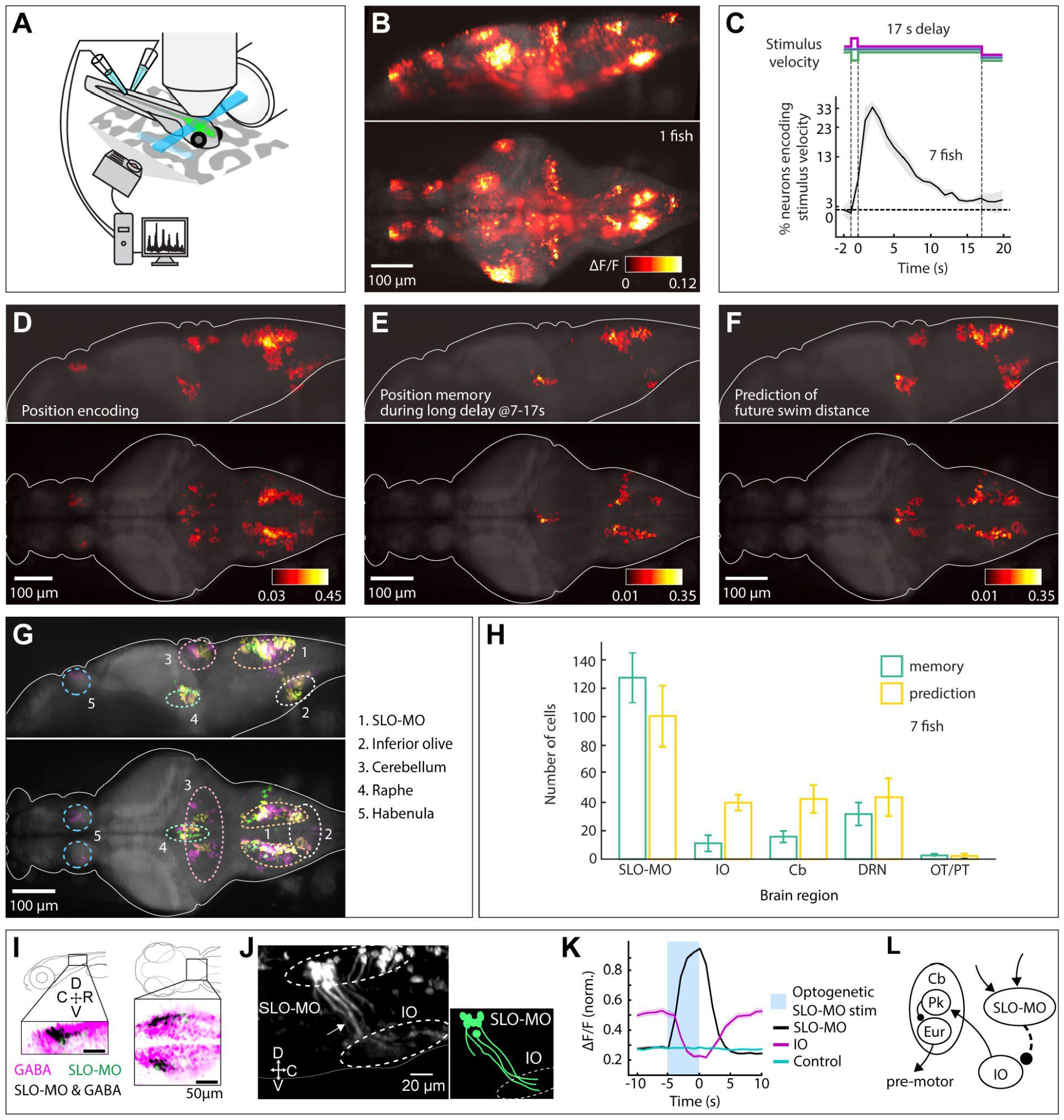
Whole-brain activity maps reveal neural population encoding self-location. (A) Virtual reality system for paralyzed zebrafish and light-sheet microscope for imaging whole-brain cellular activity. (B) Example brain-wide activity following forward visual motion, just before swim initiation (Suppl. Fig. 3 for earlier and later time points). (C) Fraction of neurons across brain activated at different time points following 1-s forward or backward translation in the virtual environment. At 17s after the shift, a fraction of neurons still encodes shift direction. (D) Whole-brain map of neurons encoding self-location during *displace, hold*, and *swim* periods. Color bar represents averaged Spearman correlation coefficient to location. Cells with p<0.005 for Spearman correlation in at least 80% of time points are shown (Methods). (E) Map of neurons encoding self-location during long *hold* period (C). Cells with p<0.005 for Spearman correlation to self-location at every time point between 7 to 17 seconds are shown. Color bar represents averaged Spearman correlation coefficient of all time points. (F) Map of neurons whose activity at *swim* period onset (before first swim bout) predicts total distance swum in *swim* period. Cells with p<0.005 for Spearman correlation at both first two time points in the *swim period* are shown. Color bar represents averaged Spearman correlation coefficient. (G) Combined maps of D-F listing brain areas where neurons occur. (H) Numbers of neurons (cell segments) (per fish) per area encoding memory of location during *displace* and *hold*, and numbers of neurons predicting future distance swum in *swim*. (I) Dorsal hindbrain map of SLO-MO neurons and GABAergic neurons in *Tg(elavl3:GCaMP6f; gad1b:RFP)* showing strong overlap between SLO-MO and GABAergic populations (black). (J) Tracing of SLO-MO neurites through sparse expression in *Tg(gad1b:Gal4; UAS:GCaMP6f)* using fluorescence correlation to SLO-MO cell body activity and anatomical tracing shows innervation of IO. (K) Optogenetic activation of SLO-MO neurons in *Tg(gad1b:Gal4; UAS:CoChR; elavl3:jRGECO1b)* shows IO neurons are inhibited during SLO-MO activation, indicating functional connection consistent with GABAergic inhibition. (L) Local circuit diagram of hypothesized (dashed line) SLO-MO inhibition of IO and known circuitry from IO to Cb cell types (Pk: Purkinje cells; Eur: eurydendroid cells, homologous to deep cerebellar nuclei).

To investigate location encoding and its transformation to behavior, we analyzed whole-brain activity of fish in virtual reality assays for (a) cells that encode positional shifts in the *displace* period (using 17s pause data as above), (b) cells whose activity (at *swim* period onset) predicts how far the fish swims in the *swim* period (with 10s pause), and (c) cells that encode fish location during the *swim* period (after 10s pause; Methods).

These analyses produced three whole-brain activity maps (Fig. 2D-F) that were consistent with each other (Fig. 2G) and can guide functional analyses and perturbation experiments to pinpoint causal mechanisms. Four spatial clusters of neurons were consistent in the maps. The most populous group, in the dorsal hindbrain, we named Self-LOcation encoding Medulla Oblongata neurons (SLO-MO; Fig. 2G, (1)). Smaller clusters were present in inferior olive (IO)^55,56^ (Fig. 2G, (2)), cerebellum^56,57^ (Fig. 2G, (3)), and dorsal raphe nucleus (DRN)^49^ (Fig. 2G, (4)). A cluster in the habenula showed activity correlating to position during swimming (Fig. 2D,G, (5)), but because this region did not encode location during the *hold* period, and correlations may be indirectly induced by swimming, we did not analyze it further.

We quantified which of these neurons encoded spatial memory or predicted future behavior. SLO-MO was biased to memory-encoding (Fig. 2H). The IO and cerebellum were biased to behavior-predicting neurons (Fig. 2H). The DRN contained both types in similar amounts (Fig. 2H). For comparison, we found neither neuron type in optic tectum (OT) or pretectum (PT) (Fig. 2H)^33,58–60^. We hypothesized that a transformation takes place between populations biased to representing spatial shifts to populations informing future swimming along a functional SLO-MO⇢IO→Cb pathway (⇢: functional connection; →: known monosynaptic connection).

To investigate SLO-MO’s cell types and functional connections, we used transgenic fish lines with GCaMP expressed in all neurons and a red co-label in cells with specified neurotransmitters. Most SLO-MO cells were GABAergic (Fig. 2I). To trace their projections, we expressed GCaMP in a sparse subset of GABAergic neurons and imaged them in the forward/backward shift assay. By correlating imaged voxels with ΔF/F signals from functionally identified SLO-MO cells, we identified projections to the area of IO dendrites (Fig. 2J). To test for functional coupling, we optogenetically stimulated functionally identified SLO-MO cells while imaging IO cells and found that IO activity was inhibited when SLO-MO was stimulated (Fig. 2K). Thus, SLO-MO, IO, and thereby cerebellum are functionally connected (Fig. 2L). We hypothesize that these regions contain the core circuit underlying representation of self-displacement and its transformation to corrective locomotion.

To investigate this transformation, we first analyzed how SLO-MO activity represents self-location in the assay of Fig. 1E: repeated *swim, hold, displace, hold, swim* periods (Fig. 3A). Two broad neuron classes contained information about fish location (Fig. 3B). Even during zero-displacement trials, many neurons had nontrivial dynamics, such as ramping up or ramping up then down, after the end of *swim* (Fig. 3C,D, *black* traces). Relative to this zero-displacement activity, activity of the first neuron class increased when fish were pulled backward (Fig. 3C), whereas activity of the second class did the opposite (Fig. 3D). Average activity over all SLO-MO cells in these classes shows that location encoding decays slowly over about 20 seconds (Fig. 3E,F). Visualized in dimensionally reduced principal component space, average population trajectories split into three branches corresponding to *backward-pull, zero-displacement*, and *forward-push* trials, which remained separated during the *hold* period, then converged during *closed-loop* period as fish trajectories converge, and returned to near the starting point (Fig. 3G; Suppl. Fig. 4A,B). Thus, representations of position are encoded within neural activity. Across nine fish (1347 SLO-MO neurons), SLO-MO cell activity correlated to fish location for over 15 seconds after displacement (Fig. 3H; Suppl. Fig. 4C). As controls, such persistent correlations were absent in motor-correlated hindbrain neurons (Fig. 3I) and pretectal neurons (Fig. 3J). Persistent effects of displacement were also visible in average activity of forward/backward tuned neural populations (Fig. 3K). SLO-MO neurons also represented location during more complex random trajectories (Fig. 3L) and other trajectory types (Suppl. Fig. 5). Cells in the SLO-MO region could encode sideways shifts (Fig. 3M), suggesting that the population code generalizes to 2D locations in space. We conclude that, across a variety of trajectory types, SLO-MO neurons persistently encode self-location.

**Figure 3:**
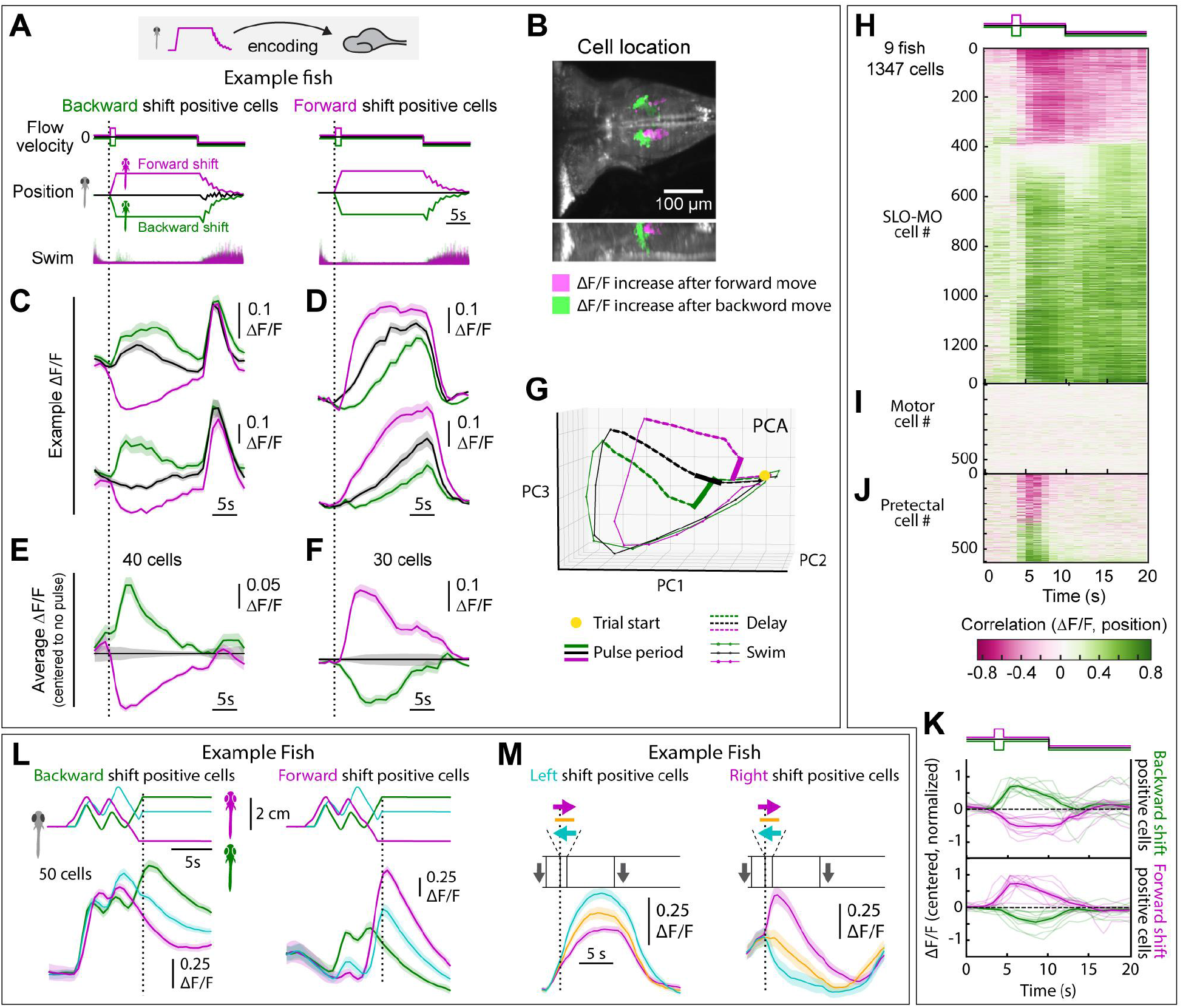
SLO-MO neuronal activity encodes self-location. (A) Encoding of self-location (relative to starting point set at end of previous *swim* period) by SLO-MO neurons during forward-push/backward-pull assay. (All fish icons are enlarged and not to scale.) (B) Location of neurons with increasing activity following backward pull (green) vs. forward push (magenta) in example fish. (C) Example neurons whose activity increases relative to zero-displacement following backward pull and decreases following forward push. (D) Example neurons with increasing activity following forward-push relative to zero-displacement activity. (E,F) Averages relative to activity during zero-displacement showing encoding of self-location across population. (Fish icons not to scale.) (G) PCA embedding of population activity. Trajectories remain separated throughout *hold* period, gradually converge during *swim* period, and return toward starting point. (H) SLO-MO neuron activity across 9 fish (1347 cell segments) sorted by Spearman correlation, showing consistent correlation to self-location. (I,J) For comparison, lack of correlation to self-location for cells with activity correlating to swim vigor, and for cells in pretectum showing visual encoding. (K) Averages of positive- and negative-correlating neurons from (H). (L) Encoding of self-location by SLO-MO neurons during complex location trajectories (example fish). (M) Encoding of sideways changes in self-location in SLO-MO neurons (example fish).

Detailed SLO-MO coding is complex (e.g., ramp and decay dynamics; Fig. 3C,D). We wondered whether a simple fixed decoder could still read out fish location. We trained a linear decoder that predicted fish location as a weighted sum of neuronal activity at the present time such that *predicted position(t-δ)* = *Σ_i_ w_i_* × *activity_i_(t)*, with *i* representing neuron index and *t* representing time (and δ=300 ms to account for slow calcium dynamics). This linear decoder (Fig. 4A) predicted fish location during random motion sequences and the *hold* period (Fig. 4B,C). Decoding fidelity decreased over time (compare Fig. 4D *left* to *right*). For comparison, a decoder trained on visual midbrain neurons performed poorly (Fig. 4E). The decoder predicted activity on individual trials, including *swim* periods (Fig. 4F), across the fish population during (Fig. 4G) and at the end of trials (Fig. 4H, 20 seconds after the initial displacement). Thus, SLO-MO activity can be used to decode fish location based on integrated optic flow.

**Figure 4:**
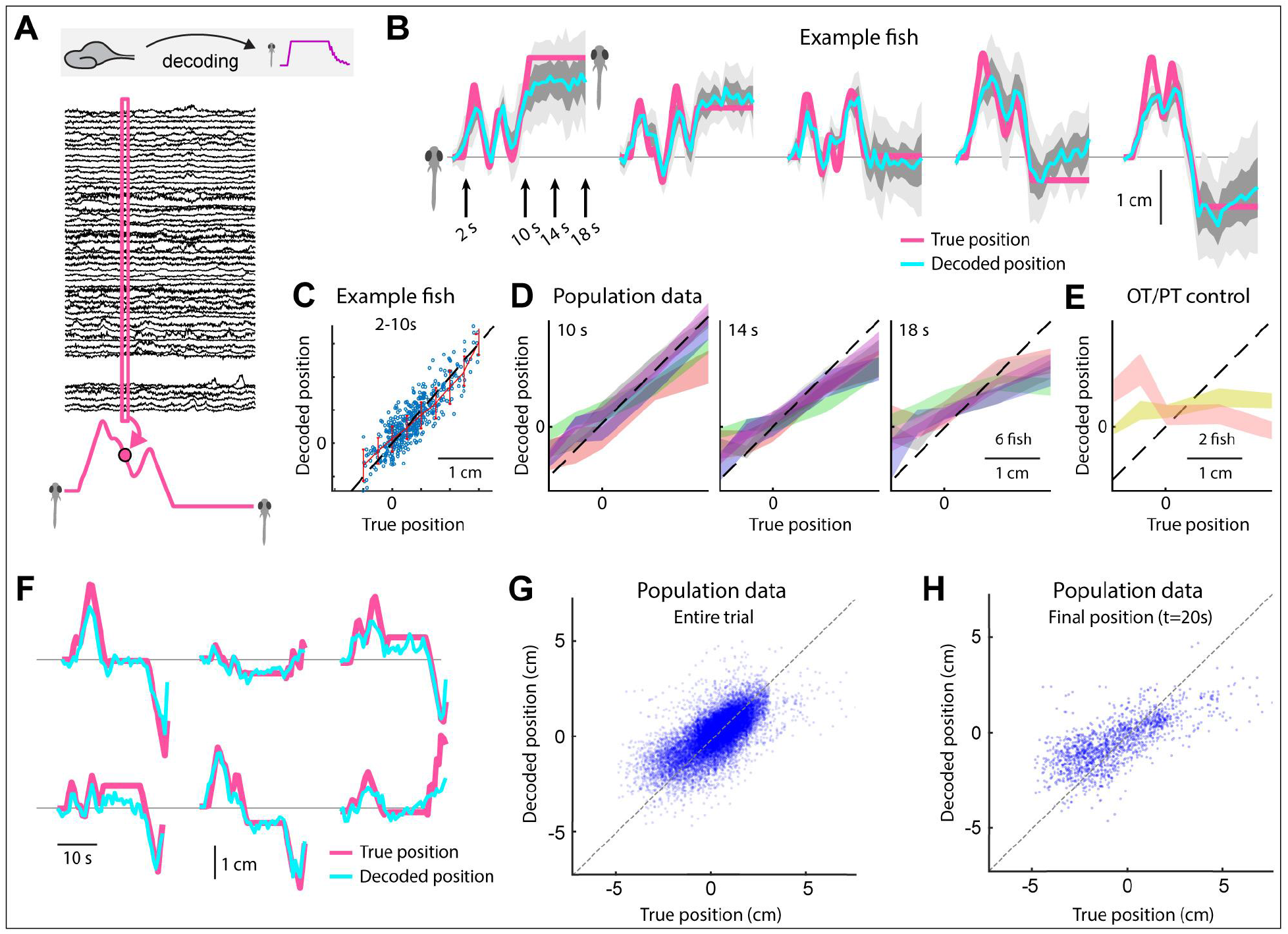
Decoding self-location from SLO-MO neuronal activity. (A) Schematic of self-location decoding from SLO-MO population activity. (B) Decoding averages of complex location trajectories in *displace* and *hold* periods of stochastic displacement assay (*dark gray*: standard error [SEM], *light gray*: standard deviation [SD]). (Fish icons enlarged, not to scale.) (C) Decoding performance throughout *displace* and *hold* periods of example fish. (D) Decoding performance at start, middle, and end of *hold* period across N=6 fish, showing gradually declining performance as SLO-MO memory decays over many seconds (colors: SEM for each fish). (E) Decoder trained on midbrain visual neurons performs poorly. (F) Example decoding single trials including first *swim* period. (G) Decoding self-location during entire trial across N=7 fish, r = 0.54, p < .01. (H) Performance of decoding self-location at end of *swim* period, r = 0.68, p < .01.

## SLO-MO is necessary for positional homeostasis and changes future locomotion

We tested whether SLO-MO neurons are necessary for the behavioral memory of position shifts, using two-photon ablation (Fig. 5A). We first imaged responses to displacements, performed online analysis to identify SLO-MO neurons (Methods^61^), then ablated all SLO-MO neurons identified as encoding a backward pull or forward push. Ablating either population also disabled the persistence of visual responses to the other (Fig. 5B), although short-timescale sensory responses remained intact, consistent with integration occurring via recurrent connectivity^62–64^. Behaviorally, post-ablation fish no longer corrected for previous positional shifts in either direction (Fig. 5C,D; Suppl. Fig. 6A-H), no matter which population was ablated. Fish still responded to visual flow (e.g., swimming in Fig. 5C), and control ablations of nearby non-SLO-MO neurons did not affect spatial memory (Fig. 5D). Thus, SLO-MO lesions abolish positional homeostasis.

**Figure 5:**
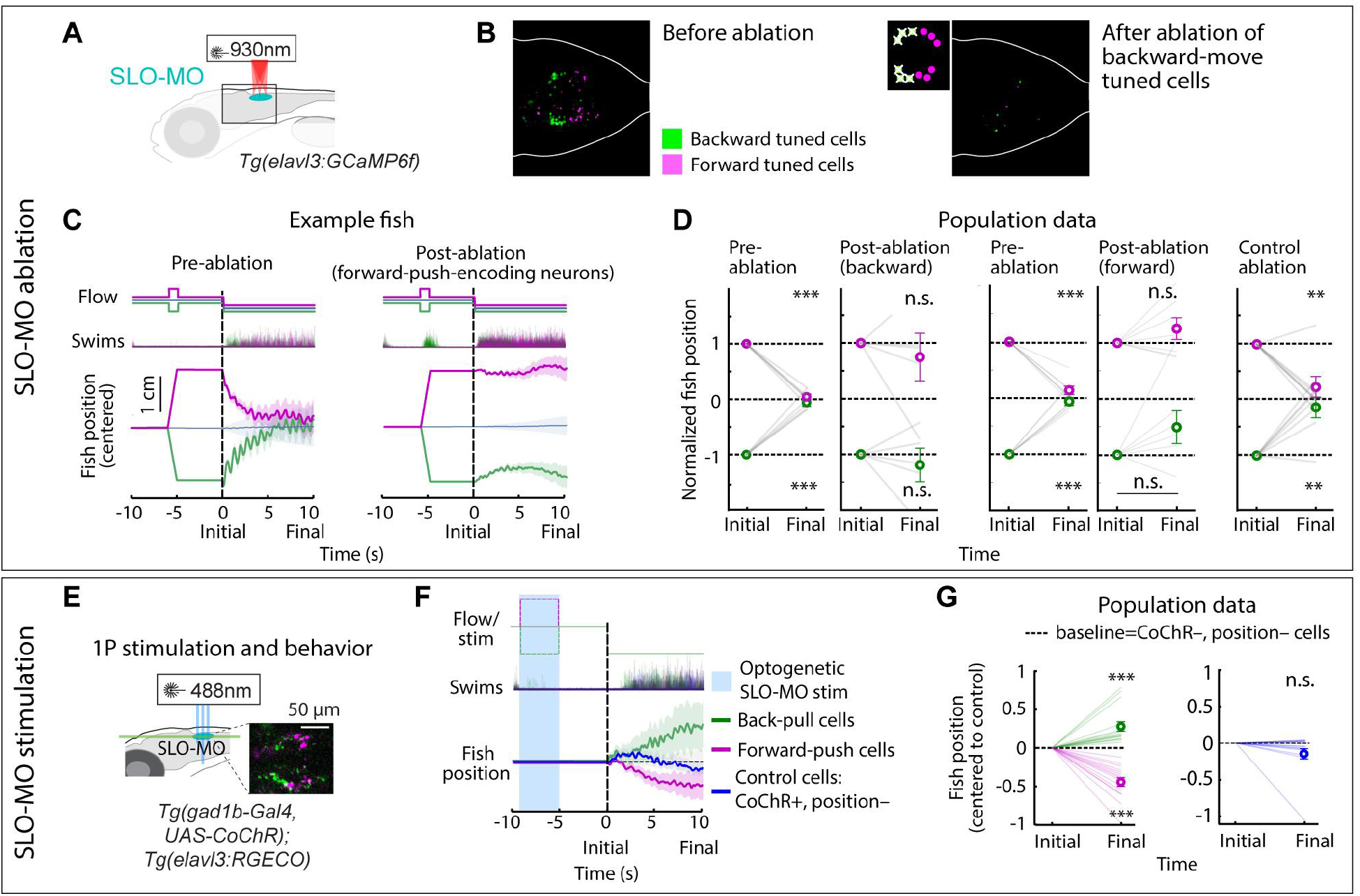
SLO-MO is necessary for location memory and changes future locomotion. (A) Two-photon laser ablation of select functionally identified SLO-MO neurons in *Tg(elavl3:GCaMP6f)* fish. (B) Ablating either population (here backward-pull encoding) abolishes position memory capacity of the entire population but leaves short-timescale sensory responses intact, suggesting recurrent connectivity within SLO-MO. (C) Ablating neurons tuned positively to forward-push abolishes positional homeostasis. (D) Population data showing consistent abolishment of positional homeostasis across N=6 fish after ablation of either forward-push or backward-pull encoding neurons, but not for nearby control neurons. (One sample t-test, ***p<0.001, p<2.5e-5 for all pre-ablation, ** p<0.01,p=3.8e-3 for backward-pull after control ablation, p=6.4e-2 for forward push after control ablation, n.s. p>0.05.) (E) Stimulation setup for activating functionally identified SLO-MO populations using optogenetics in *Tg(gad1b:Gal4; UAS:CoChR; elavl3:jRGECO1b)* fish. (F) Optogenetic activation of backward-pull encoding neurons (green) causes increased swimming 5-10s later and activation of forward-push encoding neurons (magenta) causes decreased swimming. Control cells that do not encode location do not affect swimming. (G) Population data showing consistent increases or decreases in swim distance following stimulation of backward-pull or forward-push encoding neurons. (15 fish, one-sample t-test, *** p<0.001, p=4.9e-4 for stimulation of backward-pull SLO-MO, p=2.1e-6 for stimulation of forward-push SLO-MO, n.s. p>0.05, p=0.15 for stimulation of control cells.) Control stimulation of cells not encoding location causes much smaller change in swimming (presumably a residual effect of visual responses to blue light).

To test if SLO-MO activity influences future swimming, we optogenetically activated specific sets of SLO-MO neurons. After functional identification of SLO-MO cell types, we optogenetically stimulated a subset of either forward-push-encoding or backward-pull-encoding neurons (~3-7 neurons; Fig. 5E). After a 5-second pause, fish behavior during the *swim* period was altered as if an actual spatial shift had occurred (Fig. 5F,G; Suppl. Fig. 6I-J). Control stimulations did not have such effects (Fig. 5G, *right*). Thus, activation of SLO-MO subpopulations mimics the behavioral effects of true spatial memory. Moreover, a transient optogenetic pulse leads to long-lasting behavior effects, suggesting that path integration contributes to the computation of spatial memories in SLO-MO, because there are no sustained cues that could mimic environmental landmarks. Based on this and the near-complete elimination of memory after SLO-MO ablation, we conclude that, in our behavioral assays, SLO-MO is the locus of spatial position memory.

## Inferior olive transforms SLO-MO-encoded memories into behavior

To explore the functional connection from SLO-MO to IO (Fig. 2I-L), we optogenetically stimulated functionally identified SLO-MO cells (Fig. 6A), which caused inhibition in a subset of IO neurons (Fig. 6B, same data as Fig. 1K). Stimulating nearby non-SLO-MO cells did not have this effect (Fig. 6C). We made functional hindbrain maps showing locations of inhibited cells (Fig. 6D). The IO, cerebellum, and anterior hindbrain (AHB) were the main loci where activity was inhibited. IO and cerebellum are involved in sensorimotor control^65,66^, and AHB codes sensorimotor cues relevant for optomotor behavior^28–30,67^. We thus hypothesized that modulation of IO activity by SLO-MO cells allows fish to combine memories of past positional shifts with responses to new visual flow.

**Figure 6:**
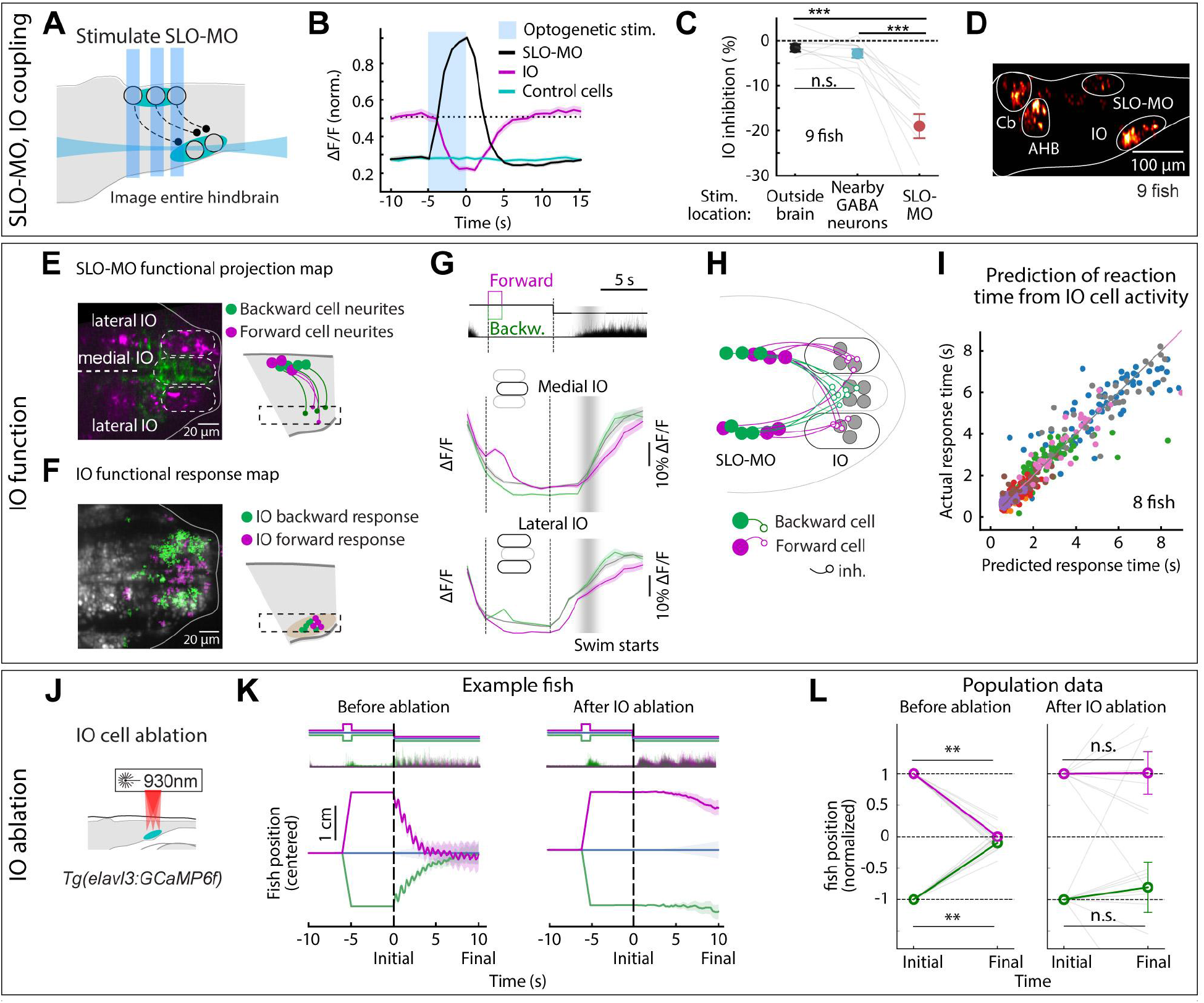
IO is functionally downstream of SLO-MO and necessary for positional homeostasis. (A) Experimental setup for stimulating functionally identified SLO-MO neurons while imaging IO neurons in fish with calcium indicator in all neurons and CoChR in mosaic subset of GABAergic neurons in *Tg(gad1b:Gal4; UAS:CoChR; elavl3:jRGECO1b)* fish. (B,C) Optogenetic stimulation of SLO-MO neurons causes inhibition of IO neurons. Stimulation of CoChR positive non-SLO-MO neurons causes no significant IO response. (p=1.7e-7 by one-way ANOVA, 9 fish. *** p<0.001, by Tukey’s post hoc test. Same data as Fig. 1K.) (D) Hindbrain map of decreases in cell activity during SLO-MO activation showing strong presence in IO, cerebellum, and anterior hindbrain (AHB). IO and cerebellum suppression is consistent with SLO-MO connections to IO. Activity decreases in AHB may be due to connections from cerebellum. Scale bar, 100 μm. (E) Imaging projections from SLO-MO in IO area in *Tg(gad1b:Gal4; UAS:GCaMP6f; elavl3:H2B-jRGECO1b)*. Green *gad1b* channel shows anatomical segregation of projections of SLO-MO neurons that encode forward-pushes (medial IO) vs. backward-shifts (lateral IO). Magenta (green) stands for forward-move-positive (backward-move-positive) responsive SLO-MO cells. (F) Imaging IO neurons (same fish as in (E); red pan-neuronal channel shown) showing segregation of neurons responding positively to forward-pushes (medial IO) vs. backward-pulls (lateral IO) — the reciprocal of (E) consistent with inhibition of IO by SLO-MO projections. Magenta (green) stands for forward-moves (backward moves) of fish in VR. (G) Time courses of IO responses to location changes. Medial IO is persistently suppressed relative to baseline following backward pulls. Lateral IO is persistently suppressed relative to baseline following forward pushes. (H) Hypothesized connectivity between SLO-MO and IO consistent with (A-H). (I) Predicting time of first swim bout during *swim* from IO activity at onset of *swim* (before first bout) across 8 fish. Swim time can be predicted, consistent with premotor role of IO. (J) Schematic of two-photon IO cell ablation. (K) Example fish positional homeostasis before IO ablation showing intact positional memory, and same fish after IO ablation showing complete loss of positional homeostasis. (L) Population data before and after IO ablation, showing consistent loss of ability to correct for positional shifts after ablation. Animals still swim in response to instantaneous flow but memory expression is lost. Thus, IO is necessary for self-location memory and positional homeostasis. (9 fish, Wilcoxon sign-rank test, ** p<0.01, p=7.6e-3.)

To probe the relationship between SLO-MO and IO activity, we imaged activity in SLO-MO projections reaching the IO area simultaneously with activity of IO neurons. We found a spatial organization of SLO-MO projections: SLO-MO activity elicited by backward pulls activated SLO-MO neurites near the medial IO, whereas SLO-MO activity elicited by forward pushes activated SLO-MO neurites near the lateral IO (Fig. 6E). The IO showed complementary responses: the medial IO was more active during forward pushes, and the lateral IO was more active during backward pulls (Fig. 6F,G), consistent with spatially organized inhibition of IO neurons by SLO-MO inputs (Fig. 6H).

Medial IO was activated when fish swam (Fig. 6G, *top*), whereas lateral IO was activated when forward visual motion began (Fig. 6G, *bottom*). Consistent with the lateral IO eliciting a positive swim drive, that region was tonically suppressed following a forward push (Fig. 6G, *bottom, magenta*) and relatively tonically elevated following a backward pull (Fig. 6G, *bottom, cyan*). Tonic suppression is consistent with persistent inhibitory input from SLO-MO. If the IO determines the behavioral response, IO activity should predict properties of future behavior. We constructed a linear predictor of response time from IO responses to forward visual motion before swimming (the first two imaging frames after stimulus onset). This model accurately predicted variability in response times from variability in IO activity (Fig. 6I). Lateral IO predicted fast responses, whereas medial IO predicted slow responses (Suppl. Fig. 7), indicating that SLO-MO-encoded memory suppresses specific IO-cerebellum sensorimotor pathways for different types of swimming.

To test the causal role of IO in future swimming, we ablated IO neurons at single-cell precision using two-photon laser ablation (Fig. 6J). Post-ablation fish were no longer able to perform positional homeostasis (Fig. 6K,L), even though they still responded to the stimulus (e.g., swim traces in Fig. 4K). Where intact fish resemble the simulated fish of Fig. 1D, *bottom*, fish with ablated IO resemble the memoryless fish of Fig. 1D, *middle*. Thus, the IO is necessary for spatial memory to influence behavior.

## Discussion

Here we report on a multiregional circuit for self-localization and positional homeostasis discovered in larval zebrafish. This system integrates visual flow to form a persistent representation of past displacement which then modulates activity in downstream circuits to influence future behavior. Because the ability to track self-location is ubiquitous amongst animal species and there are numerous homologies across vertebrate brains, it may have homologues across vertebrates, including mammals. If so, it may operate in parallel to, or eventually be read out by, hippocampal and entorhinal cortical circuits.

The visual circuitry upstream of SLO-MO is still unknown. Visual circuits responding to optic flow include the pretectum^28–30,33^ which may directly or indirectly connect to SLO-MO neurons. Likewise, the organization of connectivity between downstream olivocerebellar circuits and premotor circuits mediating the behavioral change remains to be determined. Finally, the precise circuit architecture enabling integration and persistent activity^68^ in SLO-MO is still unknown. The abolishment of persistence but preservation of short-timescale responses following ablation of part of SLO-MO suggests that recurrent connectivity within this region supports integration. Connectivity analysis through electron microscopy^69,70^ and neuron morphology atlases^71^ is likely to clarify these questions.

The DRN only encoded backward displacement (Suppl. Fig. 9); because of the near-complete effect of SLO-MO ablation on positional homeostasis, we suspect DRN’s effect is modulatory (as in short-term motor learning^49^) and that it does not directly affect positional homeostasis. However, indirect effects on the interplay between short-term motor learning and positional homeostasis must be taken into account when constructing complete circuit models for flexible fish behavior.

Integrating circuits for heading direction, eye movements, and information accumulation for decision making have been found in many species^15,16,34,72,73^. Behaviors based on path integration have also been found in multiple species ^36,38,74^; information integration underlies many computations in the brain^34^. In addition to SLO-MO, the hindbrain contains multiple circuits capable of persistent activity, such as the oculomotor integrator^75–77^. These circuits may share similar network architectures for integration^78,79^, and may be adjacent or partially overlap^61,80–83^.

In relation to mouse neuroanatomy^84^, we hypothesize that SLO-MO overlaps with the homologue of the nucleus prepositus hypoglossi (NPH)^85,86^, based on GABAergic cell type and functional connectivity to inferior olive^85,86^, as well as location relative to area A2^87^ (also known as NE-MO^48^), the nucleus of the solitary tract, and the fourth ventricle^88^ (Suppl. Fig. 8). The NPH has been studied in the context of eye movements, but it may also play a role in navigation in mammals. Modern dense electrode arrays^89^ may help uncover a navigational function of homologous hindbrain circuits in mammals.

Sideways displacements of the fish in VR also elicited population patterns of long-timescale activity in cells in the SLO-MO area. Similarly slow responses to sideways motion are found in the anterior hindbrain and pretectum^28,30^, suggesting these areas may jointly mediate the initial turn in the direction of flow before one-dimensional positional homeostasis is triggered as animals face the flow.

The functional connection of SLO-MO to the olive and, thereby, cerebellum is consistent with effects of cerebellar perturbations on hippocampal place cell activity^90^, which will be clarified by future connectivity data in multiple species.

Positional homeostasis can be thought of as minimizing, over time, a ‘displacement error’. The framework of cerebellum-mediated error signaling may link positional homeostasis and motor control. In a control theory framework, SLO-MO represents the integration term, without which the controller becomes memoryless (Fig. 1D, middle) as shown in fish behavior after SLO-MO or IO ablation (Fig. 5C,D, Fig. 6K, Suppl. Fig. 6A-H, Suppl. Fig. 7B-E). This suggests a hierarchical circuit structure in which a memoryless feedforward network mediates the basic optomotor response, modulated by olivocerebellar pathways for behavioral flexibility^91–93^, which in turn are modulated by SLO-MO pathways that introduce displacement memory to enable positional homeostasis.

Elements of cognition such as evidence accumulation and decision making^28,30,48^ and behavioral state switching^48,94–96^ have been shown to involve dynamics in unexpected areas of the zebrafish brain, including the hindbrain. Our results on positional homeostasis are consistent with the idea that more ancient brain regions also form substrates of higher-order behaviors, and increasing evidence points to the involvement of unexpected subcortical regions in higher-order mouse behaviors^97^. This emerging view that cognitive processes are widely distributed across the nervous system aligns with the evolutionary concept that complex behaviors emerged, in part, by building new circuits on top of ancient brain structures that perform related computations^98^. Brain-wide surveys of neural activity may thus be critical for determining the mechanisms of distributed cognitive function.

## Acknowledgements

This was supported by the Howard Hughes Medical Institute and by the Simons Foundation (Simons Collaboration on the Global Brain #542943SPI). We thank Albert Lee, Behtash Babadi, Eyal Gruntman, Bradley Hulse, Xinyu Zhao, Sandro Romani, Scott Sternson, David Stern, Vivek Jayaraman, and Florian Engert for discussions and comments on the manuscript, and Minoru Koyama, Paul Tillberg, Greg Fleishman, Martha Bagnall, Robert Baker, David Kleinfeld, and Lauren McElvain for discussions. We thank the GENIE Project Team at Janelia Research Campus for the jGCaMP7 plasmids. We are grateful to the Janelia Research Campus vivarium for support with transgenesis and for fish care, to Marisa Dreher for contributing to figure design, and to Janelia Experimental Technology for help with microscopy.

## Supplementary Figures

**Supplementary Figure 1.**
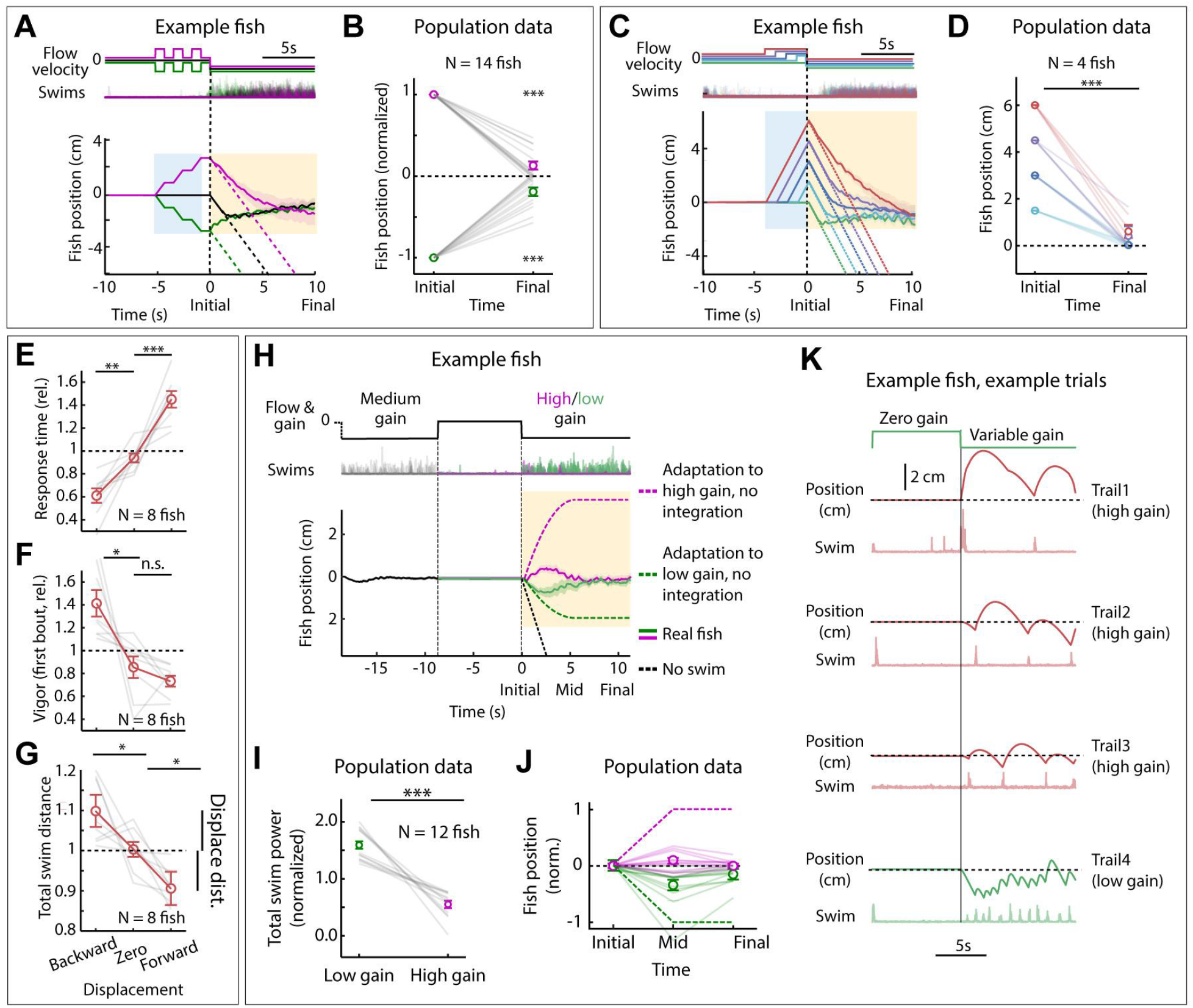
Positional homeostasis during complex trajectories and motosensory gain changes. (A) Triple forward push/backward pull assay. Example fish, showing converging trajectories, indicating successful integration across three consecutive forced displacements. (B) Population data (14 fish) showing near-complete correction for earlier displacement. (One sample t-test, *** p<0.001, p=9.7e-10 for forward push, p=1.9e-10 for backward pull.) (C) Assay to test integration over varying durations. An example fish successfully integrates displacement and corrects for it in the *swim* period. (D) Population data (4 fish) showing accurate correction, i.e. path integration. (Two-tailed paired t-test, ***p<0.001, p=1.08e-7 for all forward push) (E) Animals respond faster (slower) after a backward pull (forward push), consistent with Fig.1 F. (Two-tailed paired t-test, **p<0.01, p=1.76e-3 for backward pull, *** p<0.001, p=3.1e-4 for forward push.) (F) Animals respond more vigorously after a backyard pull, consistent with Fig.1 F. (Two-tailed paired t-test, * p<0.05, p=0.0151 for backward pull, n.s. p>0.05, p=0.115 for forward push). (G) Total swim distance (normalized) corresponds to the earlier displacement, consistent with Fig.1 F. (Two-tailed paired t-test, * p<0.05, p=0.031 for backward pull, and p=0.012 for forward push). (H) After motosensory gain changes (high: ×2, low: ×0.5), animals still integrate position. *Dashed lines*: position of model fish performing gain adaptation (linear adjustment of vigor over 5 seconds) but no path integration. Solid lines: position of real fish. (I) Swim vigor during low or high motosensory gain shows gain adaptation. (Two-tailed paired t-test, *** p<0.001, p=1.6e-5.) (J) Average fish position in low vs. high gain trails initially diverges, then converges. *Dashed lines*: normalization to model fish performing gain adaptation but no path integration. (K) Example trials during high and low motosensory gain (data from E-G) showing accurate positional homeostasis in both cases.

**Supplementary Figure 2.**
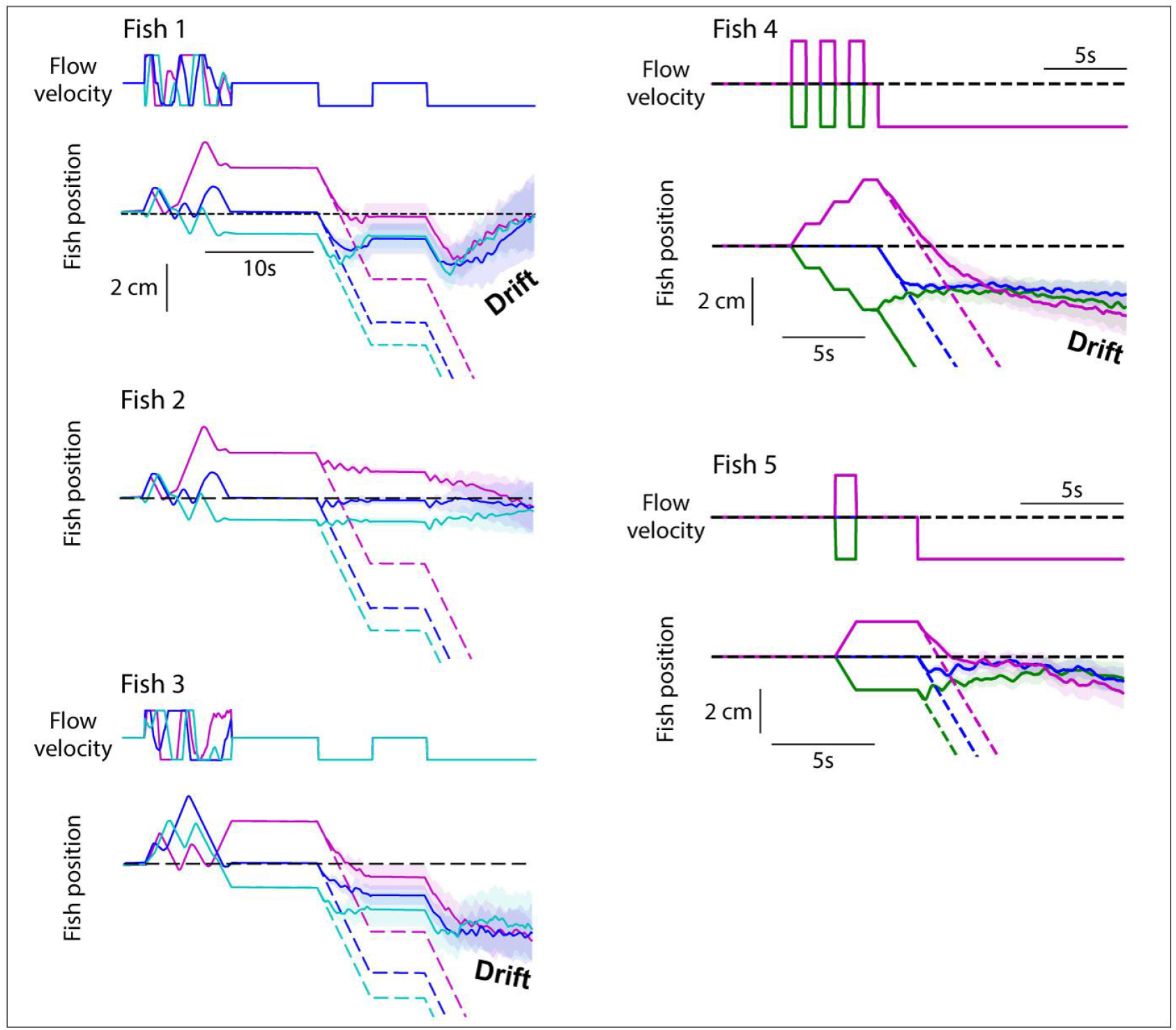
Positional drift accumulates over time. Example fish in various patterns of forced displacements, showing forward drift (fish 1) and backward drift (fish 3, 4) and no or negligible drift (fish 2, 5). Drift tends to be larger when the fish is being imaged with the light sheet switched on, possibly reflecting input noise acting on the sensory velocity signal.

**Supplementary Figure 3.**
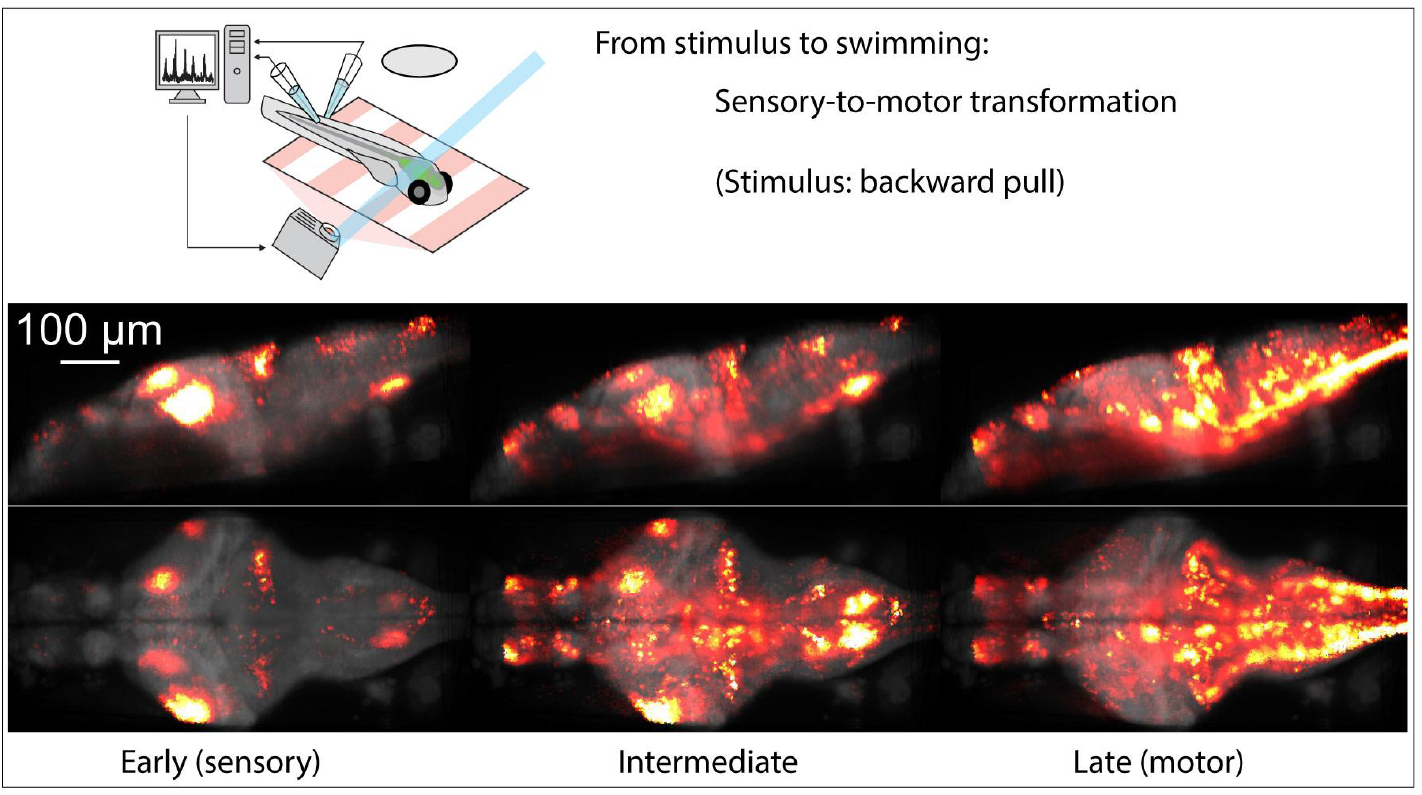
Whole-brain activity during a sensorimotor transformation. Brain activity averaged over trials in an example fish. *Left*: in the 300 ms following onset of a backward pull through VR (forward visual motion from the point of view of the fish). *Middle*: at an intermediate time, after stimulus onset, just before (0-300 ms) swim onset. *Right*: immediately following (0-300 ms) swim onset.

**Supplementary Figure 4.**
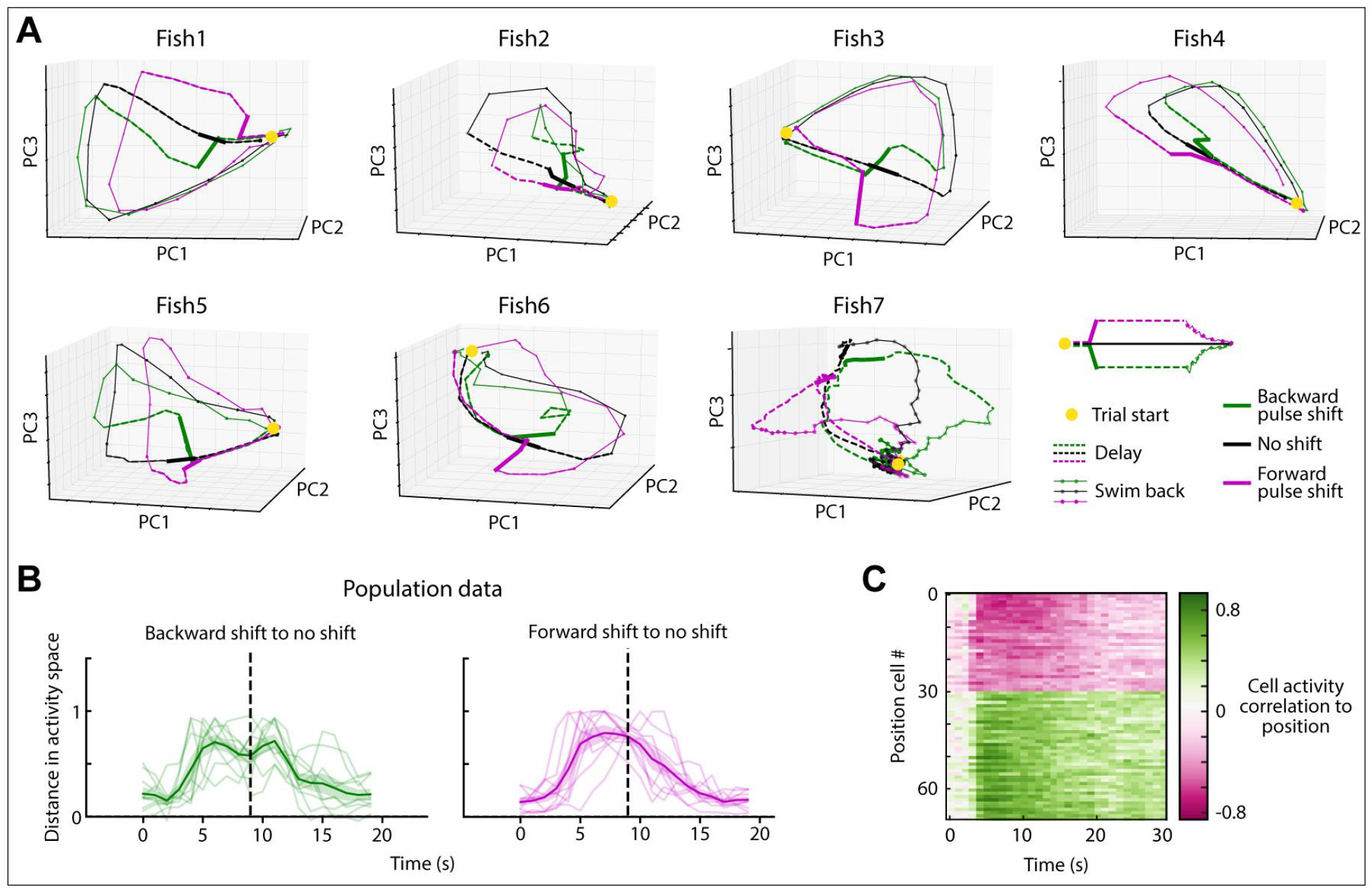
Network trajectories for individual fish, and related data. (A) Trajectories through dimensionally-reduced network space for seven individual fish, related to Fig. 3. Dimensionality reduction done with Principal Component Analysis. (B) Distance in PC space (first 6 components) between backward-shift and zero-shift, and backward-shift and no-shift activity, for 9 fish. Since distances are zero or greater, noise causes distance to always be positive even before the spatial shift at ~2 seconds. (C) Ranked correlation of cell activity to fish position for example fish of Fig. 3A-G, also related to population data of Fig. 3H.

**Supplementary Figure 5.**
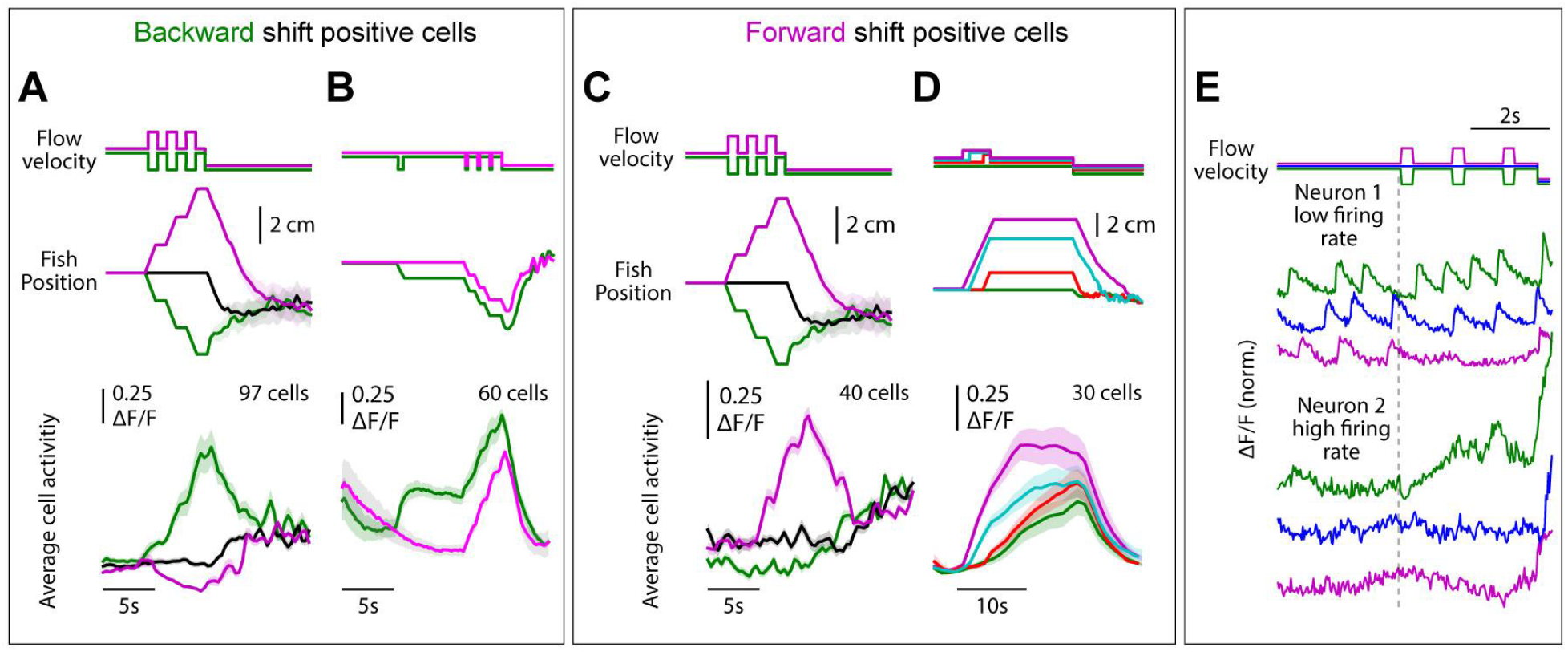
SLO-MO activity during various complex trajectories. (A) Neurons (in an example fish) with increasing activity following three consecutive backward or forward shifts, showing integration over the shifts. (B) Neurons (in an example fish) showing integration over backward shifts. The *green* trials are offset by one additional shift 5 seconds before the triplet shifts; this is reflected in a persistent increase in neuronal activity. (C) Integration across triplet shifts by neurons that respond positively to forward shifts. (D) Additional example of integration by SLO-MO neurons during forward shifts of varying durations followed by *hold* periods. (E) Individual neuron traces showing integration across triplet shifts, fast imaging of single planes at 33 Hz. The firing rate of neuron 1 is low enough that individual spikes are visible in the calcium trace, showing persistently increasing spike rate after backward shifts and persistently decreasing spike rate after forward shifts. The firing rate of neuron 2 appears to be higher and, although individual spikes are not visible due to sampling rate and calcium indicator limitations, firing rate can be seen to increase following a backward shift relative to a forward shift.

**Suppl. Figure 6.**
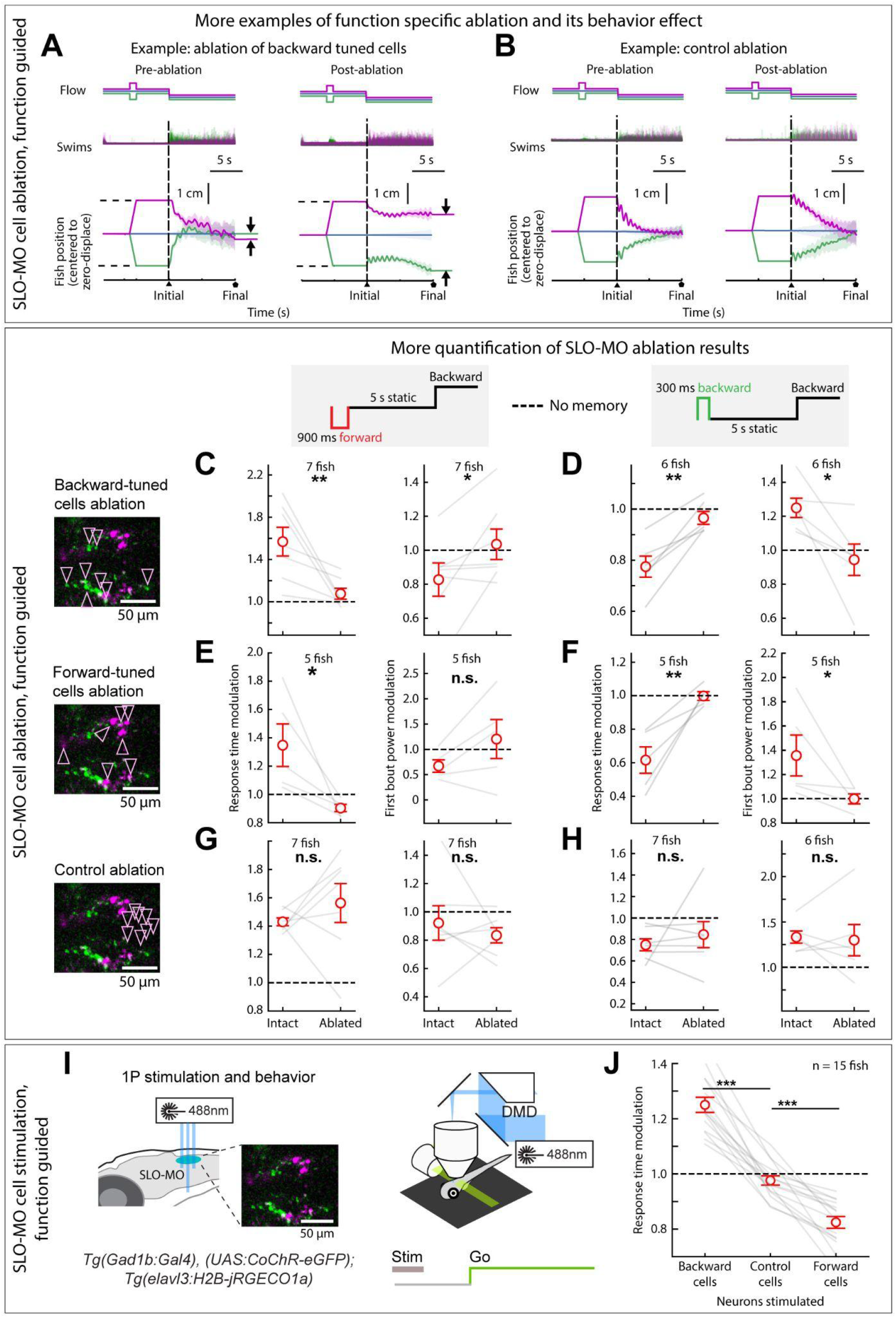
Detailed characterization of effects of SLO-MO ablation and stimulation. (A) Example fish showing effect of ablation of backward-pull-tuned cells, to complement example fish shown in Fig. 5C. (B) Example of effect of control ablation (non-SLO-MO cells in the dorsal hindbrain), showing intact ability to correct for past displacements. (C-H) Additional quantification of the effects of SLO-MO ablation. C,D: ablation of backward-tuned cells. E,F: ablation of forward tuned cells. G,H: control ablation of nearby non-SLO-MO neurons. (Two-tailed paired t-test, ** p<0.01, * p<0.05, n.s. p>0.05.) (I) Diagram of optogenetic stimulation and functional imaging setup. (J) Optogenetic excitation of functionally-identified SLO-MO neurons causes changes in the time to first swim bout in the *swim* period, consistent with changes in distance swum shown in Fig. 5F,G. (Two-tailed paired t-test, ***p<0.001)

**Supplementary Figure 7.**
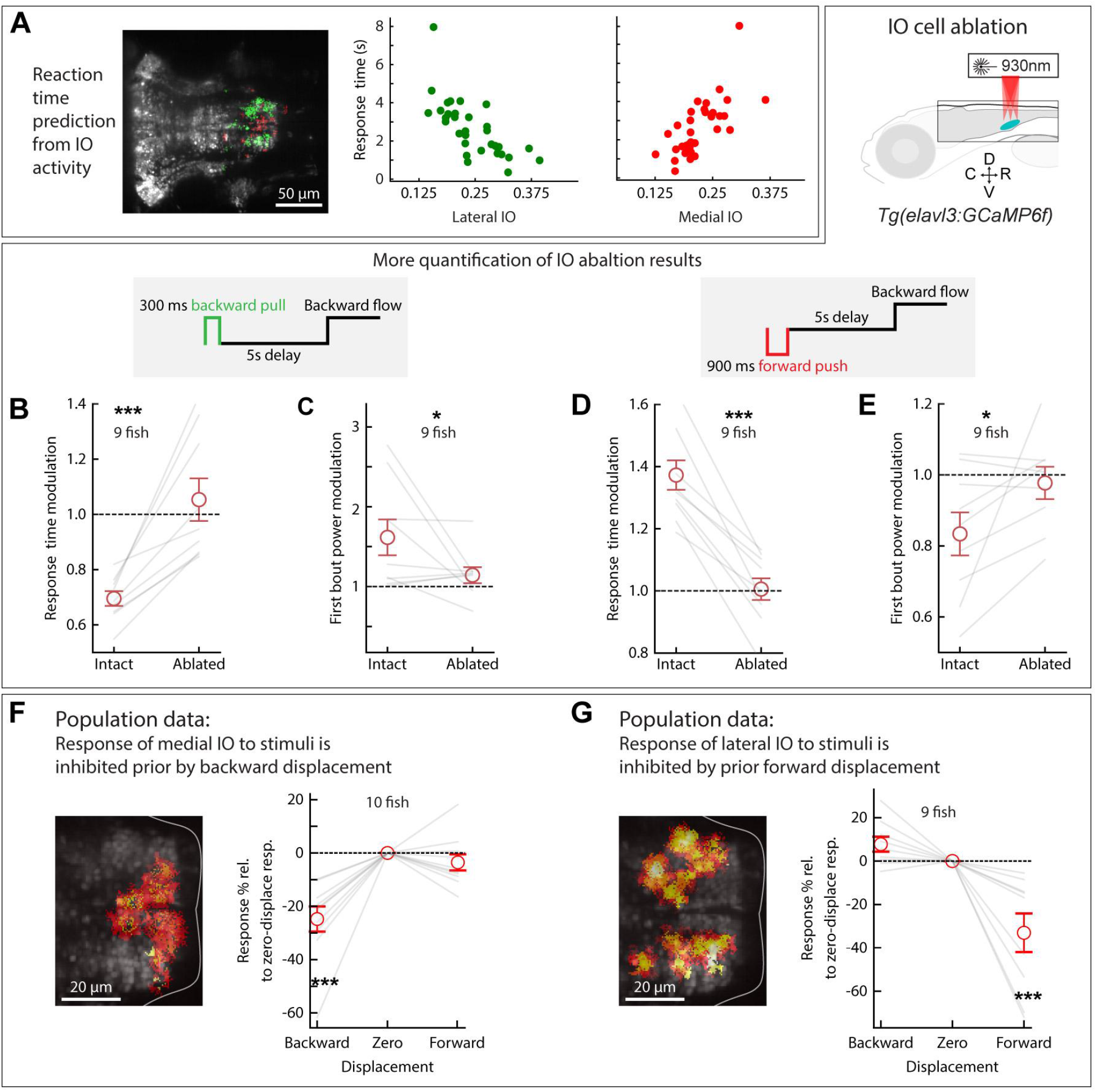
Inferior Olive dynamics, and effects of IO ablation on behavior. (A) Correlation between lateral and medial IO activity and reaction time, complementing the joint decoder of Fig. 6I. (B-E) Additional quantification of the effects of IO lesions on response time and swim power. (9 fish, two-tailed paired t-test, *** p<0.001, * p<0.05.) (F-G) Backward displacements suppress medial IO activity; forward displacements suppress lateral IO activity. Population data to complement Fig. 6G. (One sample t-test, *** p<0.001.)

**Supplementary Figure 8.**
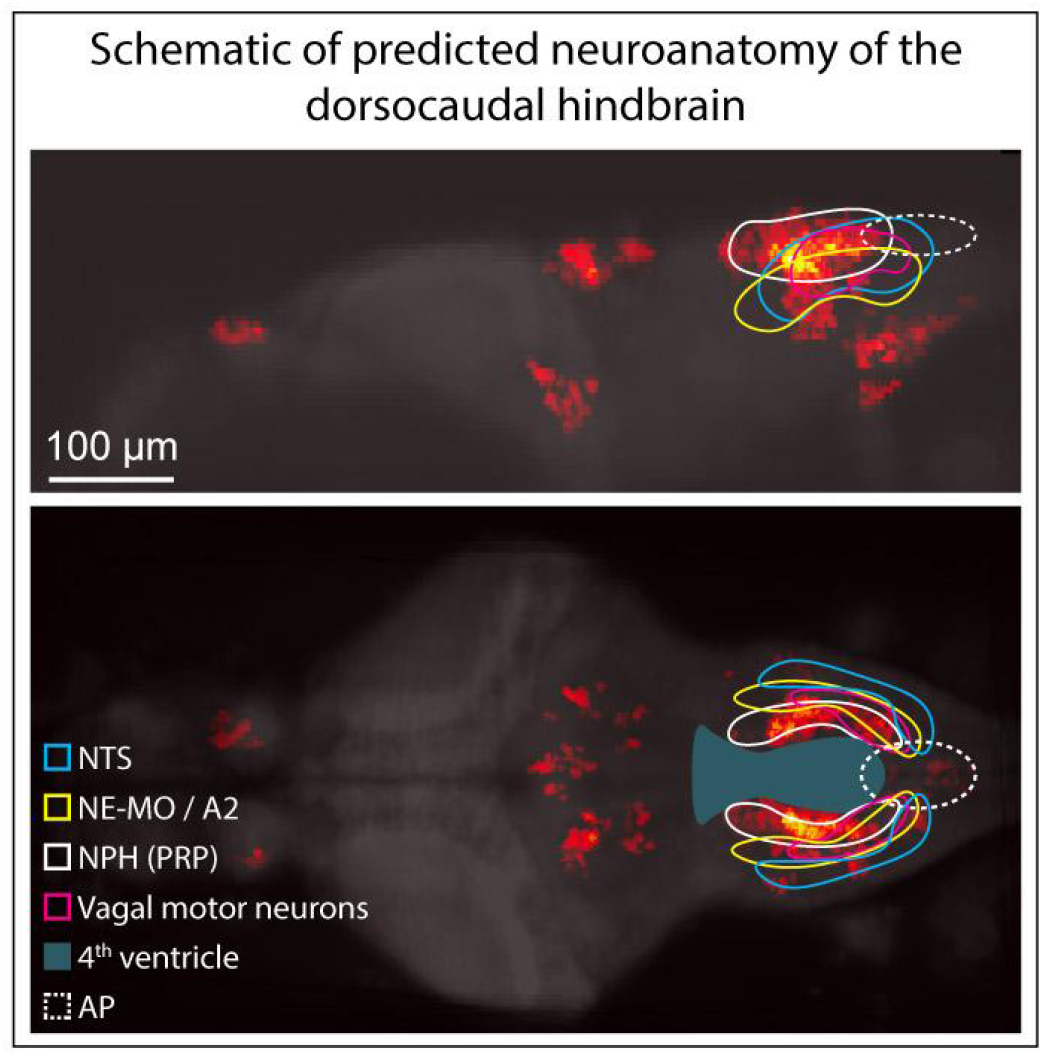
Known and hypothesized neuroanatomy of the dorsocaudal larval zebrafish hindbrain. This schematic, based on multiple zebrafish and mouse publications and atlases, accompanies the hypothesis that SLO-MO overlaps with the homologue of the nucleus prepositus hypoglossi (NPH), a major GABAergic input to the IO^85,86^. SLO-MO neurons are also GABAergic and project to and inhibit the IO. Our results are consistent with a monosynaptic connection between SLO-MO and IO, and we showed that activating SLO-MO functionally inhibits IO. SLO-MO neurons are spatially arranged relative to the nucleus of the solitary tract (NTS), noradrenergic area A2^87^ (called NE-MO in^48^), and are close to the fourth ventricle (Suppl. Fig. 1H) (Tabor et al. 2019); the NPH also has these properties (Allen Mouse Brain Atlas, http://www.brain-map.org,^84^). Historically, the NPH has been studied in the context of eye movement control, but (parts of) the NPH may be multifunctional. Circuits for positional homeostasis and for oculomotor control have in common that they must integrate visual slip into a persistent signal^75–77^, so it would not be surprising if some sharing or overlap between these circuits was present, possibly with similar mechanisms for persistent activity.

**Supplementary Figure 9.**
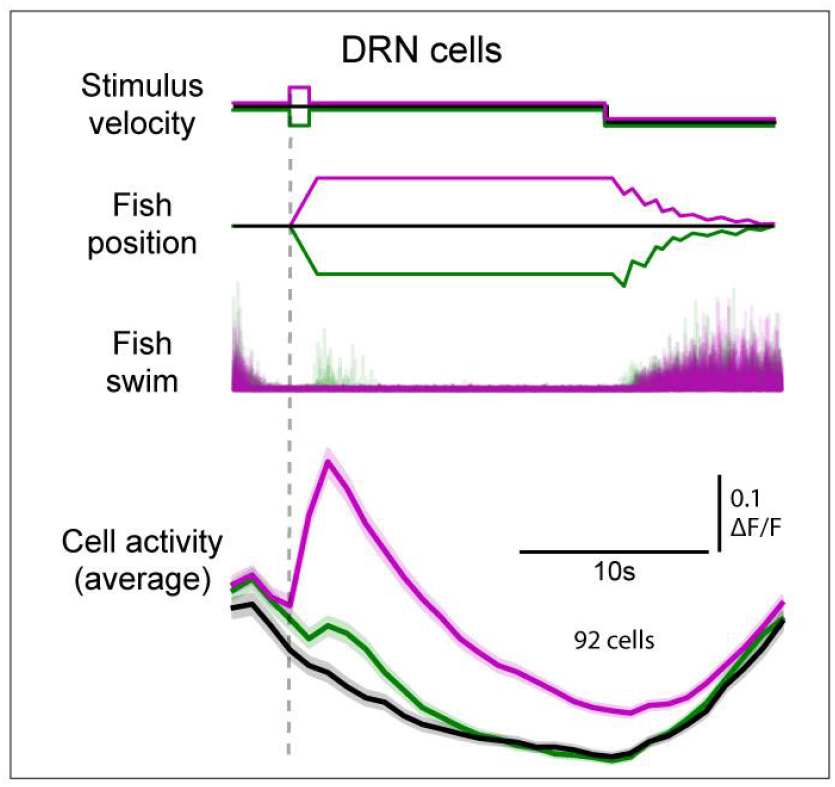
Dorsal Raphe neurons persistently encode forward pushes but not backward pulls. Activity of a subset of dorsal raphe neurons increases persistently when fish are pushed forward (*magenta*), but no average persistent change relative to zero-displacement was found (*green*, compare to *black*).

## Methods

(*for A brainstem integrator for self-localization and positional homeostasis - En Yang, Maarten F. Zwart, Mikail Rubinov, Ben James, Ziqiang Wei, Sujatha Narayan, Nikita Vladimirov, Brett D. Mensh, James E. Fitzgerald, Misha B. Ahrens*)

### Experiments and fish care

All experiments presented in this study were conducted according to the animal research guidelines from NIH and were approved by the Institutional Animal Care and Use Committee and Institutional Biosafety Committee of Janelia Research Campus. All larvae were reared at 14:10 light-dark cycles according to standard protocols at 28.5°C (Westerfield 2000). Zebrafish from 5 to 7 dpf were fed rotifers and used for experiments. Zebrafish sex cannot be determined until ~3 weeks post-fertilization (Uchida et al. 2002), so experimental animals’ sex was unknown.

### Transgenesis

Transgenic zebrafish larvae were in casper or nacre background (White et al. 2008).

*TgBAC(glyt2:loxP-DsRed-loxP-GFP)* (Satou, Kimura, and Higashijima 2012), *TgBAC(slc17a6b:loxP-DsRed-loxP-GFP)* (Satou et al. 2013) and *Tg(gad1b:loxP-RFP-loxP-Gal4)^jf99^* were all used without Cre-mediated recombination and referred to as *Tg(glyt2:dsRed), Tg(vGlut2a:dsRed)* and *Tg(gad1b:dsRed)* respectively.

*Tg(elavl3:GCaMP6f)^jf1^, Tg(elavl3:H2B-GCaMP6f)^jf7^, Tg(elavl3:ReaChR-TagRFP-T)^jf10^*, and *Tg(elavl3:jRGECO1b)^jf17^* lines are described (Dana et al. 2016; Dunn et al. 2016; Vladimirov et al. 2018). *Tg(Gad1b:Gal4)^jf49^, (UAS:GCaMP6f)^jf46^* and *Tg(gad1b:Gal4)^jf49^, (UAS:CoChR-eGFP)^jf44^* are described in (Mu et al. 2019).

*Tg(elavl3:jGCaMP7f)^jf96^*, *Tg(elavl3:H2B-jGCaMP7f)^jf90^*, *Tg(elavl3:H2B-jRGECO1a)^jf112^* lines were generated with Tol2 system (Urasaki, Asakawa, and Kawakami 2008) and published jGCaMP7f and jRGECO1a constructs (Dana et al. 2016, 2019).

For imaging of neural activity we used transgenic zebrafish lines expressing:

- Cytosolic GCaMP6f, *Tg(elavl3:GCaMP6f)^jf1^* (Dunn et al. 2016)
- Cytosolic jGCaMP7f, *Tg(elavl3:jGCaMP7f)^jf96^* (this paper)
- Nuclear-localized GCaMP6f, *Tg(elavl3:H2B-GCaMP6f)^jf7^* (Dunn et al. 2016)
- Nuclear-localized jGCaMP7f, *Tg(elavl3:H2B-jGCaMP7f)^jf90^* (this paper)
- Cytosolic jRGECO, *Tg(elavl3:jRGECO1b)^jf17^* (Dana et al. 2016)
- *Tg(Gad1b:Gal4)^jf49^;Tg(UAS:GCaMP6f)^jf46^* (Mu et al. 2019)
- *Tg(Gad1b:Gal4)^jf49^;Tg(UAS:jGCaMP6f)^jf46^;Tg(elavl3:H2B-jRGECO1a)^jf112^* (this paper)

For neurotransmitter identity determination we used:

- *Tg(vGlut2a:dsRed)* - (*TgBAC(slc17a6b:loxP-DsRed-loxP-GFP)*, (Satou et al. 2013))
- *Tg(glyt2:dsRed)* - (*TgBAC(glyt2:loxP-DsRed-loxP-GFP, (Satou, Kimura, and Higashijima 2012)*)
- *Tg(gad1b:dsRed)* - (*Tg(gad1b:loxP-RFP-DsRed-loxP-Gal4)^jf99^)* (Kler et al. 2021)

For optogenetic stimulations we used:

- *Tg(gad1b:Gal4)^jf49^; Tg(UAS:CoChR-eGFP)^jf44^* (Mu et al. 2019)
- *Tg(gad1b:Gal4)^jf49^; Tg(UAS:CoChR-eGFP)^jf44^; Tg(elavl3:H2B-jRGECO1a)^jf112^* (this paper)
- *Tg(elavl3:ReaChR-TagRFP-T)^jf10^* (Dunn et al. 2016)

### Light-sheet microscopy

The design of our light-sheet microscope and behavior setup is as described earlier (Vladimirov et al., 2014, 2018). Briefly, the detection arm consisted of a water-dipping detection objective (16x/0.8 NA, Nikon) mounted vertically on a piezo stage (Physik Instrumente), a tube lens and an sCMOS camera (Orca Flash 4.0, Hamamatsu). The detection arm was equipped with a band-pass filter (525/50 nm, Semrock) and long-pass filter (590 nm, Semrock) for separating GCaMP and RGECO fluorescence light from scattered 488 nm and 561 nm laser light, respectively. Each of the two illumination arms consisted of an air illumination objective (4x/0.28 NA, Olympus) mounted horizontally on a piezo stage, a tube lens, an f-theta lens and a pair of galvanometer scanners (Cambridge Technology). The scan mirrors and f-theta lens scanned the collimated laser beam (488 nm / 561 nm) laterally and along the z-axis of the image space. The excitation laser was rapidly turned off electronically every time the laser was scanned over the eye of the fish to avoid direct stimulation of the retina.

### Preparation of zebrafish for fictive behavior and imaging experiments

All animal handling procedures were done as previously described (Vladimirov et al. 2018; Kawashima et al. 2016). Larval zebrafish at 6-7 dpf were paralyzed by immersion in 1 mg/ml alpha-bungarotoxin solution (Invitrogen) dissolved in external solution (in mM: 134 NaCl, 2.9 KCl, 2.1 CaCl_2_, 1.2 MgCl_2_, 10 HEPES, 10 glucose [pH 7.8]; 290 mOsm). Duration of immersion was empirically determined, ranging from 20 to 40 seconds. Once paralyzed, the fish were embedded using 2% low-melting point agarose (Sigma-Aldrich Inc.) in a custom made chamber (designs available on request), with agarose removed from the animal to expose the tail (for electrophysiological recordings (Ahrens et al. 2012)) and head (for imaging). Extracellular recordings from the tail were made with borosillicate pipettes (TW150-3, World Precision Instruments) pulled by a vertical puller (PC-10, Narishige) and shaped by a microforge (MF-900, Narishige) to have a tip diameter of approximately 40 micrometers. Electrodes were filled with collagenase (1% w/v; Sigma-Aldrich Inc.) dissolved in fish rearing water. Recordings were made using an Axon Multiclamp 700B amplifier in current clamp setting and acquired using National Instruments DAQ boards. Signals were sampled at 6 kHz, and band-pass filtered with a 3 kHz/100 Hz low-pass/high-pass cutoff. To identify swim bouts, signals were further processed by taking the local standard deviation in a sliding 10 ms window, after which a simple thresholding operation was used to identify swims. Signals were recorded and visual stimuli presented using custom software written in C# (Microsoft; (Ahrens et al., 2012)). A diffusive plastic screen was attached to the bottom of the chamber from the outside to allow image projections from below using a miniature projector (Sony Pico Mobile Projector; MP-CL1) connected to the behavior acquisition and control computer. Experiments were performed in 1D virtual environments consisting either of finely spaced (~2 mm thickness) red/black bars or irregular random red/black patterns to image GCaMP fish; green/black gratings or patterns were used when imaging jRGECO1b fish. Irregular patterns (Fig. 1C) were generated by creating a 2D Gaussian random phase noise and weighting it in Fourier space with a weighting function proportional to 1/(freq_x_ + freq_y_), inverse Fourier transforming it, scaling it to lie between 0 and 1, and thresholding it at 0.32 and 0.68 to create noise patterns as in Fig. 1C. For the optogenetics experiment we used dim blue-black gratings with identical parameters to prevent nonspecific activation of ReaChR channels; all three colors reliably elicited the OMR.

### Imaging of neural activity

Whole brain imaging using light-sheet microscopy as in Figure 2 was performed using both light-sheet illumination arms on volumes spanning 300 μm along the dorso-ventral axis (61 z-planes, 5 μm intervals) and 820 and 410 μm along the longitudinal and lateral axes in *Tg(elavl3:jGCaMP7f)^jf1^* fish; imaging of the hindbrain in Figures 3–6 was performed using only the lateral illumination arm on volumes with variable dimensions spanning the regions of interest, ranging from 400 to 600 μm along the longitudinal axis. High-speed calcium imaging of SLO-MO neurons in Suppl. Fig. 5E, was performed using *Tg(Gad1b:Gal4; UAS:GCaMP6f)* by imaging a single plane around the SLO-MO cluster at 33 Hz in the lightsheet microscope. High-speed calcium imaging of IO neurons in Fig. 6E and F, Suppl. Fig. 7A, was performed using *Tg(Gad1b:Gal4)^jf49^;Tg(UAS:jGCaMP6f)^jf46^;Tg(elavl3:H2B-jRGECO1a)^jf112^* by imaging 9 planes (45 μm thick in total) around IO clusters.

### Behavioral assays

Either red/black gratings of spatial period 0.4 cm, or red/green textured pattern were presented to the fish, alternating between periods of backward, stationary, and forward motion. In all behavioral experiments, response times were determined by detecting the first swim after the start of the *swim* period. Bout power was defined as the integrated windowed standard deviation of the electrical signal from the tail recording as in (Dunn et al. 2016).

### Visual stimulus delivery

A fish holder was mounted on the transparent acrylic bottom section of the chamber, which simplified the process of removing agarose surrounding the animal. A diffusive plastic screen was placed ~1 cm below the fish allowing image projections from below using a miniature projector emitting red monochromatic images. The visual stimulus was generated using a Sony Pico Mobile Projector (MPCL1) connected to the behavior acquisition and control computer.

### Neuron ablations

To ablate individual neurons we used two-photon laser plasma ablation based on published protocols and optimized to act on sub-single-neuron spatial scales (Fang-Yen et al. 2012; Vogel and Venugopalan 2003). We used high power laser (500 mW, measured after the objective) at 930 nm wavelength with a short exposure time (1-2 ms for neurons 0-100 μm from the dorsal brain surface and up to 3-4 ms in ventral areas 150-200 μm deep) using a Coherent Chameleon Ultra II laser. The time interval between successive ablation points was set to 5 seconds to allow heat and chemicals released by cell ablation to dissipate and minimize unintended brain damage. The choice of short exposure and long inter-ablation interval allowed to avoid cavitation (water vapor bubbles) in the brain tissue. Longer exposure times tended to produce off-target effects whereby such bubbles occurred, and single-neuron resolution was lost; restricting the exposure time to 1-4 ms avoided such effects and produced sub-neuron-scale cell damage.

In the behavioral tests, the optomotor (OMR) assay was started 20 minutes before ablation, followed by an ablation session lasting 5-10 minutes, followed by further OMR behavior trials for another 30-40 minutes after ablation.

### Optogenetics

For the optogenetic stimulation of SLO-MO neurons while imaging calcium transients across the hindbrain, we used a digital micromirror device (DMD)-based (Lightcrafter discovery kit, Texas Instruments) method of delivering 488 nm light pulses to predefined ROIs with approximately single-cell precision (for more details, see (Mu et al. 2019)). *Tg(gad1b:Gal4)^jf49^; Tg(UAS:CoChR-eGFP)^jf44^; Tg(elavl3:H2B-jRGECO1a)^jf112^* fish at 6 d.p.f. were imaged using a 561 nm excitation laser in a 10 minute experiment in which the OMR was induced by alternating 10-second periods of stationary and forward moving gratings with a brief pulse inserted during stationary grating, as described in Figure 1. SLO-MO neurons were identified using pixel wise Spearman correlation after cross correlation registration (Freeman et al. 2014; Miri et al. 2011), with maps cross-referenced to those showing CoChR expression patterns in each fish. In each experiment, 5 to 10 CoChR-expressing SLO-MO neurons were selected for subsequent stimulation, with the same number of nearby non-SLO-MO CoChR-expressing cells and an area outside the brain selected as controls.

For Fig. 6A-D, the 488 nm stimulation laser was pulsed at 10 Hz with 60 ms pulses during whole-hindbrain imaging. Experiments lasted 60 minutes. Stimulation epochs were interspersed by rest periods of 40 s.

For Fig. 5E-G, the same setup and DMD stimulation paradigm (10 Hz, 60 ms pulses) were used. Stimulation epochs were interspersed by rest periods of 40 s. The behavioral paradigm consisted of 10 s *hold* alternating with 10 s *swim* as described in the main text.

### Extraction of cells from voxel data

We extracted populations of cell bodies and sections of neuropil (‘cell segments’) from raw fluorescence data with a volumetric segmentation pipeline first described in Mu et al. (Mu et al. 2019) and available from https://github.com/mikarubi/voluseg. We first registered all volumes to the volume recorded halfway through the experiment using ANTs (Avants et al. 2010). We then defined an intensity-based brain mask and divided the masked volume into about 2000 spatially contiguous and slightly overlapping volumetric blocks. Each block contained about 100 tightly packed 5 μm diameter cell spheres.

We initialized the spatial footprint of each cell as a local-intensity peak of strongly correlated and contiguous voxels. We then used constrained non-negative matrix factorization to segment each block in parallel. For *n* voxels, *t* timepoints, and *c* cell segments we factorized,

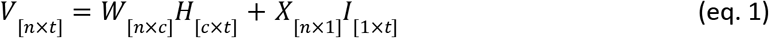

where *V* is the full spatiotemporal fluorescence matrix for each block *W* and *H* are, respectively, the spatial footprint and temporal trace of cell segments, and *X* and *I* are the corresponding spatial footprint and temporal trace of the background signal.

We solved equation 1 approximately using alternating least-squares (Berry et al. 2007). We regularized the spatial cell footprint *W* by restricting it to a contiguous segment within a 10 μm diameter sphere, and by projecting to a sparse subspace (Hoyer 2004). We compute the baseline of the resulting fluorescence with a sliding-window percentile filter that estimated the 10th percentile of the data in 5-minute windows. We then computed Δ*F/F* as

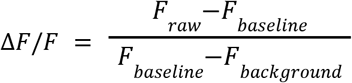

### Registration of brains onto a reference brain

To compare results across analyses we registered all experiments to a representative brain volume, using iterative rigid-body, affine and diffeomorphic non-rigid image registration, implemented in ANTs (Avants et al. 2010).

### Computation of the brain activity maps

We computed the functional maps from the time-varying jGCAMP7f intensity signal in cells segmented from methods described above. The time series of all cells segmented in the brain (segmented neurons) were first separated into multiple repeated task periods (trials). A new Δ*F/F* array of size [*l*(number of neurons segmented)×*m*(number of trials)×*t*(number of time points in each trial)] was constructed. Another array of [*m*(number of trials)×*t*(number of time points in each trial)] containing information on presented stimuli (such as past location displacement) or behavioral aspects (such as current fish body location), of the matching trials was constructed. A Spearman correlation between the two matrices was performed for all individual cells at each time point across trials, resulting in a correlation coefficient matrix and the corresponding p-value.

Fig. 2D was computed from the trial structure presented in Fig. 1E, with a 5-second hold period after the forced position displacements and a 10 seconds swim period. In order to locate neurons encoding self-positions during the entire trial including both the 5s *hold* period after displacement and the 10s *swim* period. The Spearman correlation was performed for every neuron in the brain between a [*m×t*] matrix of Δ*F/F* and a [*m×t*] matrix of current fish location (where m is the number of trials, t is the number of time points in each trial). A correlation coefficient array with length t was then computed for each individual neuron segmented, as well as its *p*-value. In this map (Fig. 2D), only neurons with persistent correlation to its current self-location are included. The selection criteria is for all time points after position displacement, at least 80% of all time points (12 seconds out of 15 seconds) having *p*<0.005. The correlation coefficient for each cell shown in the heat map is the average correlation coefficient of that cell for all the time points considered. This process was repeated for each individual fish, and all cells meeting these criteria were extracted and used for subsequent analysis.

Fig. 2E was computed from the trial structure presented in Fig. 3A, with a long *hold* period (17 seconds) after the forced position displacements in order to locate long time constant position displacement memory cells, emphasizing the delay period without behavior feedback. The Spearman correlation was performed for every neuron in the brain between a [*m×t*] matrix of Δ*F/F* and a [m×t] matrix of past forced displacement, a correlation coefficient array with length *t* was then computed for each individual neuron, as well as its p-value. In this map, only neurons with persistently correlation to the pervious displacement (*p*<0.005 during the entire last 10 seconds before *swim*, which is from 7s to 17s) are included. The correlation coefficient for each cell shown in the heat map is the average correlation coefficient of the 10s considered for that cell. This process was repeated for each individual fish, and all cells meeting these criteria were extracted and used for subsequent analysis.

Fig. 2F was computed from all fish from the trial structure presented in Fig1. E, emphasizing the go cue response of all neurons before the fish starts to swim in order to locate cells encoding information predicting swim distance for the coming trial. A new Δ*F/F* array of [*l*(number of neurons)×m(number of trials)*×2*(first two time points after go cue before fish swims)] were constructed. Another array of [m(number of trials)×2] containing information of the swim distance of the current trial was constructed. The Spearman correlation was then computed for every neuron in the brain between a [*m×2*] matrix of Δ*F/F* and a [*m×2*] matrix of future swim distances. The purpose of duplication of swim distance here (×2) is to match dimensionalities of that of the Δ*F/F* array. A correlation coefficient array with length 2 was then computed for each individual neuron, as well as its p-value. In this map, only neurons with *p*<0.005 for both time points were included. This process was repeated for each individual fish, and all cells meeting these criteria were extracted and used for subsequent analysis.

To combine data across fish, the maps obtained above from individual fish were processed via a statistical test across fish described in (Marques et al. 2020) to produce the final maps shown in Fig. 2 D-F. This statistical test ascribes a *p*-value to each neuron based on the distribution of similar functionally-defined neurons in other fish. This p-value was used to discard neurons that had no consistent nearby partners in other fish: only neurons with *p*<0.001 were kept. In this shuffle test, the null hypothesis is that the averaged minimal distance of each neuron in each individual fish to neurons of the same function class in other fish is the same as its minimal distance to a random neuron (5000 times shuffle repetition) in other fish.

The python code of the data processing pipeline is available at (https://github.com/nvladimus/ZebrafishFunctionalMaps_LinearRegression) and uses the Thunder package which can be downloaded at http://thunder-project.org (Freeman et al. 2014).

### Statistical analysis

Data were tested for statistical significance across fish by, as indicated in the text and figures:

- Two-tailed paired t-test (Suppl. Fig. 1D,E,F,G,I, Suppl. Fig. 6C-H,J, Suppl. Fig. 7B-E)
- One-way ANOVA followed by Tukey’s post-hoc test (Fig. 6C)
- Wilcoxon rank-sum test (Fig. 6L)
- One sample t-test (Fig. 1F,H, Fig. 5D,G, Suppl. Fig.1B, Suppl. Fig. 7F,G)
- shuffle test (Fig. 2D,E,F)
- Pearson regression (Fig. 4G,H)

Error bars in figures represent SEM unless otherwise stated. All the statistical analyses were performed using custom-written scripts in Python. Distribution of all population data first went through a normality test, Wilcoxon rank-sum test was used if the data was not normally distributed.

### Fish simulations

To simulate a fish’s behavior in various conditions (Fig. 1D), we utilize a PID Controller, a basic element in Control Theory (Graf 2016), although without a derivative element. The probability of a swim bout at time t is:

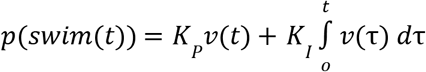

where *p* was additionally thresholded to lie between 0 and 1 and where *K_P_* and *K_I_* are constants controlling the effect of velocity and position, respectively and *v*(*t*) is the velocity of the grating including both open-loop and closed-loop, self-induced motion. In the unresponsive model fish (Fig. 1D, top), both parameters are set to zero, as no feedback drives swim bouts. In fish with no spatial memory (Fig. 1D, middle) the value of *K_P_* during the *swim* period was set to 0.022 and the value of *K_I_* to zero, reflecting the fact that only the fish’s apparent velocity, and not the memory of prior position, effects the swim drive, while remained zero. In the fish with spatial memory (Fig. 1D, bottom), the value of *K_I_* was set to 0.01 and the value of *K_I_* to 0.014, reflecting the fish’s memory of its previous *pre-displace* location. In all conditions, the displacement per swim bout was 0.8, with a minimum inter-bout interval of 1 second (30 simulated time units) used in order to approximate the fish’s natural swimming behavior. Note that here increasing the value of *K_I_* decreases the time until each *displace* condition converges, but not their final convergence. Ignoring the refractory period, the fish without spatial memory can also be viewed as a homogeneous Poisson process with parameter with constant parameter λ = *K_P_v*, while the fish with spatial memory will be recognized as a nonhomogeneous Poisson process allowing for a memory of the spatial position.

### Fish location decoding from SLO-MO activity

In order to decode a fish’s virtual position from its brain activity in response to a pseudo-random motion stimuli encompassing both open-loop (where the fish was not in control of its apparent motion) as well as closed-loop (where the fish actively controlled its apparent location) portions, we utilized a ridge regression with a leave-one-out cross validation (LOOCV) paradigm. For each trial, we set the fish’s position to zero prior to the beginning of the open-loop *displace* period and integrated the velocity for both open-loop *displace, hold*, and closed-loop *swim* periods prior to down-sampling the position to the acquisition rate of the brain imaging data, yielding position traces as seen in Fig. 4F. Each trial was then sequentially excluded from a training set, where we fit a linear decoder to the equation:

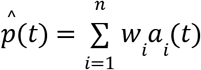

where *w* is the parameter weight, *a(t)* is the activity at time *t*, and *i* is a subscript for each neuron. To avoid overfitting and issues with multicollinearity, we regularized the fit using a ridge regression, where the regularization constant λ was linearly spaced from 0.5 to 50 in steps of 0.5. Following training for each fold, the excluded trial was then utilized for cross-validation, and the RMSE was computed between the testing trials’ true and estimated positions for all values of λ. The value of λ minimizing the average RMSE across all trials was then selected independently across fish.

### Motosensory gain adaptation simulation

To simulate behavior with motosensory gain adaptation but without positional integration (Suppl. Fig. 1H), we assumed that fish would linearly adjust swim vigor over a period of 5 seconds from [original swim vigor] to [original swim vigor]×[original motosensory gain]/[new motosensory gain]. For example, if the original swim vigor is 1 and the original motosensory gain is 1 but then changes to a new motosensory gain of 2, the model assumes that fish will linearly shift its swim vigor from 1 to 0.5 over a period of 5 seconds. This produces a parabolic position trajectory (because [fish velocity] = [drift velocity] + [swim vigor]×[motosensory gain]) as shown in Suppl. Fig. 1H.

